# Electrogenetic signaling and information propagation for controlling microbial consortia via programmed lysis

**DOI:** 10.1101/2022.12.05.519192

**Authors:** Eric VanArsdale, Ali Navid, Monica J. Chu, Tiffany M. Halvorsen, Gregory F. Payne, Yongqin Jiao, William E. Bentley, Mimi C. Yung

## Abstract

To probe signal propagation and genetic actuation in microbial consortia, we have coopted the components of both redox and quorum sensing (QS) signaling into a communication network for guiding composition by “programming” cell lysis. Here, we use an electrode to generate hydrogen peroxide as a redox cue that determines consortia composition. The oxidative stress regulon of *Escherichia coli*, OxyR, is employed to receive and transform this signal into a QS signal that coordinates the lysis of a subpopulation of cells. We examine a suite of information transfer modalities including “monoculture” and “transmitter-receiver” models, as well as a series of genetic circuits that introduce time-delays for altering information relay, thereby expanding design space. A simple mathematical model aids in developing communication schemes that accommodate the transient nature of redox signals and the “collective” attributes of QS signals. We suggest this platform methodology will be useful in understanding and controlling synthetic microbial consortia for a variety of applications, including biomanufacturing and biocontainment.

## Introduction

Synthetic microbiomes and cell consortia offer the opportunity to distribute tasks among their constituent species (Roell et al., 2019; Zhou et al., 2015). For green manufacturing, such schemes rely on both optimization of individual strains (e.g., for metabolite flux (Xu et al., 2020), substrate utilization (Akdemir et al., 2022), and replication of ecological niches (Shahab et al., 2020)) and consortia composition. Synthetic microbiomes can also inform on species functions within natural environments and guide intervention or consortia reconstruction (Clark et al., 2021). Designer consortia of varied purpose and makeup continually emerge (Johnston et al., 2020; Li et al., 2022; Rodríguez Amor & Dal Bello, 2019; Stephens et al., 2019b; van der Hoek & Borodina, 2020).

Our work is motivated by the complex environments of the rhizosphere (Scheme 1), where redox signaling cues microbial function and a symphony of signaling preserves interkingdom symbiosis, including nutrient uptake, alarmone signaling, and even pathogen abatement (Faure et al., 2009; Jo et al., 2022). Here, plant-derived hydrogen peroxide (H_2_O_2_) initiates a complex communication network that embraces direct electron transport and reaction as well as molecular signal generation, transport, interpretation, and propagation. Communication signals are sometimes short-lived, they travel both short and long distances, and there are vastly different thresholds for recognition and actuation. Some cues, such as quorum sensing (QS) autoinducer-1 (AI-1), gather cells to coordinate their function as a collective (Fuller et al., 2017; Hense et al., 2007; Nunan et al., 2020; Redfield, 2002; Teplitski et al., 2011). Others, like H_2_O_2_, are rapidly degraded and have multiple functions, including altering redox properties of a microenvironment and also activating gene expression inside cells, as it freely passes cell membranes.

**Scheme 1.**
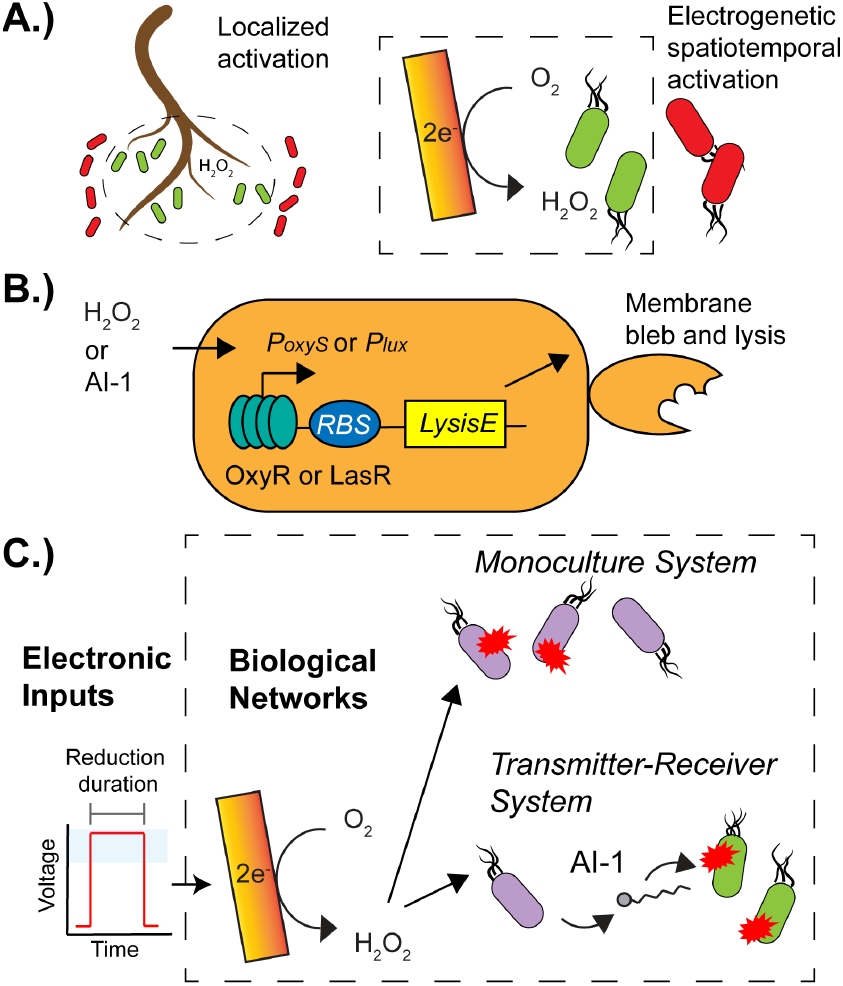
Methods for programming electrogenetic lysis into bacterial cells. **A.)** Localized H_2_O_2_ signaling within the rhizosphere results in only a small portion of microbial cells near the root becoming induced in response to a temporary signal, thereby creating a heterogenous distribution of gene activation across the total consortium. This spatiotemporally confined signaling can be mimicked using electrochemical production of H_2_O_2_ because the signal is confined to the near vicinity of the electrode and H_2_O_2_ is rapidly degraded. As a result, only cells that pass within the close vicinity of the electrode are induced into an “active” state and the proportion of H_2_O_2_ activated cells becomes dependent both on the time duration of the input signal and the signal’s voltage. **B.)** As models for microbial lysis, we programmed cells to respond to either H_2_O_2_ through the OxyR transcription factor or autoinducer-1 (AI-1) through the LasR transcription factor to regulate expression of Lysis protein E from φX174 phage (*lysisE*) which induces cell lysis. **C.)** Two separate systems were used to study signal propagation: (i) a Monoculture system in which electrode-produced H_2_O_2_ directly induces OxyR-regulated expression of LysisE, and (ii) a Transmitter-Receiver system in which H_2_O_2_ induces the synthesis of AI-1 in a “transmitter” population, which in turn regulates expression of LysisE in a “receiver” population. Synthetic biology designs and charge-dependent control over electrogenetic relay are investigated.

To gain a mechanistic understanding of the complex signaling in redox environments, we created a well-controlled experimental mimic of the native signaling processes that are initiated by H_2_O_2_ (Scheme 1A). We used a simple gold electrode, immersed in a cell-containing medium, to generate H_2_O_2_ for the transmission of a “program” into a mini-consortium. In Scheme 1b, we link the signals H_2_O_2_ and AI-1 to the expression of the gene for phage φX174 lysis protein E, *lysisE*, causing cellular lysis. Then (Scheme 1C), we explore two communication modes to learn how communication and synthetic biology designs enable understanding and application.

## Materials and Methods

### Materials, Plasmids, and Cloning

All chemicals, unless specified, were purchased from Sigma Aldrich (St. Louis, Missouri). *Escherichia coli* DH5α (Invitrogen, Carlsbad, CA) and NEB10β (New England Biolabs (NEB), Ipswich, MA) were the primary strains used in this work. *E. coli* cells were grown in Luria-Bertani (LB) medium or a minimal medium as described previously (Gustavsson et al., 2011). All electrochemistry equipment was purchased from CH Instruments (Austin, TX) except reference electrodes, which were purchased from Bioanalytical Systems, Inc. (West Lafayette, IN).

OxyR-LasI-LAA was cloned previously (Stephens et al., 2019a). K31 and K33 variants of the - LasI-LAA transmitter plasmid were prepared by PCR amplification of gene blocks containing *P*_*katG*_*-rbs31* and *P*_*katG*_*-rbs33* from Integrated DNA Technologies (IDT, Coralsville, IA) and insertion into the OxyR-LasI-LAA plasmid between the open reading frame containing *oxyR* and the transcriptional start site of the *lasI(LAA)* gene using the Golden Gate assembly method. All lysis and fluorescent reporter plasmids used in this work were cloned using IDT gene blocks and InFusion cloning (Takara Bio, San Jose, CA). A table of all plasmids and relevant features can be found in Table S1. All plasmid sequences are available upon request.

### H_2_O_2_ and AI-1 induced lysis assays

Cell lysis was evaluated by measuring the optical density at 600 nm (OD_600_) of cell cultures after either the addition of N-3-oxo-dodecanoyl-L-homoserine lactone (AI-1, Cayman Chemicals, Ann Arbor, MI), hydrogen peroxide, or electrogenetic induction via the ORR reaction (see below) in a 96-well black walled plate (Corning Inc., Corning, NY). Prior to induction, cells were first inoculated overnight in LB (37°C, 250 rpm) and then re-inoculated at a ratio of 1:50 in minimal medium. Once the cells reached mid-logarithmic growth, the cells were then diluted into either monocultures or cocultures with a fixed initial total OD_600_ of 0.1 and induced with indicated inducer concentrations. Transmitter-receiver cocultures were inoculated with the same initial total OD_600_ of 0.1 containing AI-1 producing transmitters fixed at 1% (i.e., OD_600_ 0.001). Measurement and culture post-induction were done in a TECAN Spark plate reader (Männedorf, Switzerland) pre-warmed to 37°C with linear shaking in a humidity-controlled chamber. All experiments were performed in triplicates unless otherwise indicated. Results are represented as the mean of the samples and all error bars indicate standard deviation from the mean.

### Oxygen-reduction reaction (ORR) electrogenetic induction

Cells were induced by the electrogenetic method previously described (VanArsdale et al., 2022). Briefly, catalytic formation of hydrogen peroxide was controlled by biasing a 2-mm gold disk electrode paired with a 1-cm long platinum wire counter electrode and an Ag/AgCl reference electrode. The 3-electrode electrochemical cell was fabricated using a 17 mm diameter glass vial with a fitted cap containing holes drilled into it to secure the positioning of each electrode. A small stir bar was added to each chamber to increase aeration during the ORR reaction. Electrodes were first cleaned with 7:3 piranha solution. Prior to induction, a “passivation treatment” was performed on each electrode to minimize error from repetitive use in which electrodes were biased to -0.5 V for 1800 seconds in fresh minimal medium. Cellular induction was performed by adding 1 mL of cell solution into the chamber and applying a square-wave voltage waveform between - 0.1 V to -0.7 V for durations of 0-1800 seconds (referred to as the “reduction durations”). The cell solution was then recovered from the chamber and kept in Eppendorf tubes until induction procedures were complete. The cells were then placed into a 96-well plate for measurement as detailed above. To measure electrode-generated H_2_O_2_ without cells, 1 mL of minimal medium was placed in the chamber above and square-wave voltage waveforms were applied as above. Hydrogen peroxide content was determined using a Pierce quantitative hydrogen peroxide assay (Thermo Fisher; Waltham, MA). Charge values were calculated as the integral of the measured current values with respect to time. The integrals were computed using the midpoint rule with a Δt = 0.001 seconds.

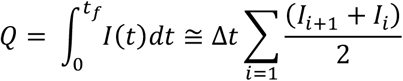

### Lysis onset and Point of Inflection calculations

For LasR and anti-ECF/ECF delay systems, the “lysis onset” was calculated as the point at which the mean of the experimental samples deviated from the mean of the null by more than 0.1 OD_600_. This threshold was chosen as it represented a departure from the null greater than the initial seed density and was greater than 10% of the final cell density in all cases. The “point of inflection” was calculated as the point at which the derivative of a 3^rd^ order polynomial fit turned negative, indicating a decrease in cell population. In mathematical models, the derivative was taken directly without a fit. Mathematical model methods and equations are in Supplemental methods.

## Results

### Redox-enabled Electrogenetic Induction of Cell Lysis

To examine cellular signaling in a “monoculture” (Figure 1), we placed *lysisE* under the control of the oxidative stress response transcription factor, OxyR (Seo et al., 2015), which upregulates, via promoters *P*_*oxyS*_ or *P*_*katG*_, many defense genes including *katG*, which encodes a bifunctional catalase-peroxidase. *P*_*katG*_ is a weaker promoter than *P*_*oxyS*_ (Barshishat et al., 2018), and its native ribosome binding site (RBS) has been engineered to provide an array of responses to different H_2_O_2_ concentrations (Rubens et al., 2016). Here, *lysisE* was cloned downstream of several previously described promoter-RBS constructs (Rubens et al., 2016), enabling a genetic screen for H_2_O_2_-induced cell lysis, where activity was scored simply as a decrease in optical density at 600 nm (OD_600_) relative to an uninduced control, instead of more a detailed analysis such as scanning electron microscopy (SEM, Figure S1). In Figure 1A, we found lysis constructs with *P*_*katG*_ with either RBS31 (red, denoted K31-LysisE) or RBS33 (purple, denoted K33-LysisE) provided a range of lysis phenotypes between 0-200 μM H_2_O_2_. RBS31 is relatively stronger than RBS33 (Rubens et al., 2016), which is reflected in increased sensitivity to H_2_O_2_ for the corresponding lysis constructs. Note that in many cases, the apparent decrease or cessation in growth was often marked by eventual regrowth, presumably owing to restoration of cell health and subsequent propagation. Constructs with the canonical *P*_*oxyS*_ promoter failed to produce colonies upon transformation, suggesting leaky expression of *lysisE* from *P*_*oxyS*_ was too toxic to the cells (data not shown). Since K31-LysisE was observed to reduce OD_600_ nearly immediately upon H_2_O_2_ addition and remained low for over 4 h, it was selected for follow-on studies. Finally in Fig 1A, we have overlayed results from a simple first-principles mathematical model (Supplemental methods) for the K31-LysisE construct showing excellent agreement with experiments.

**Figure 1.**
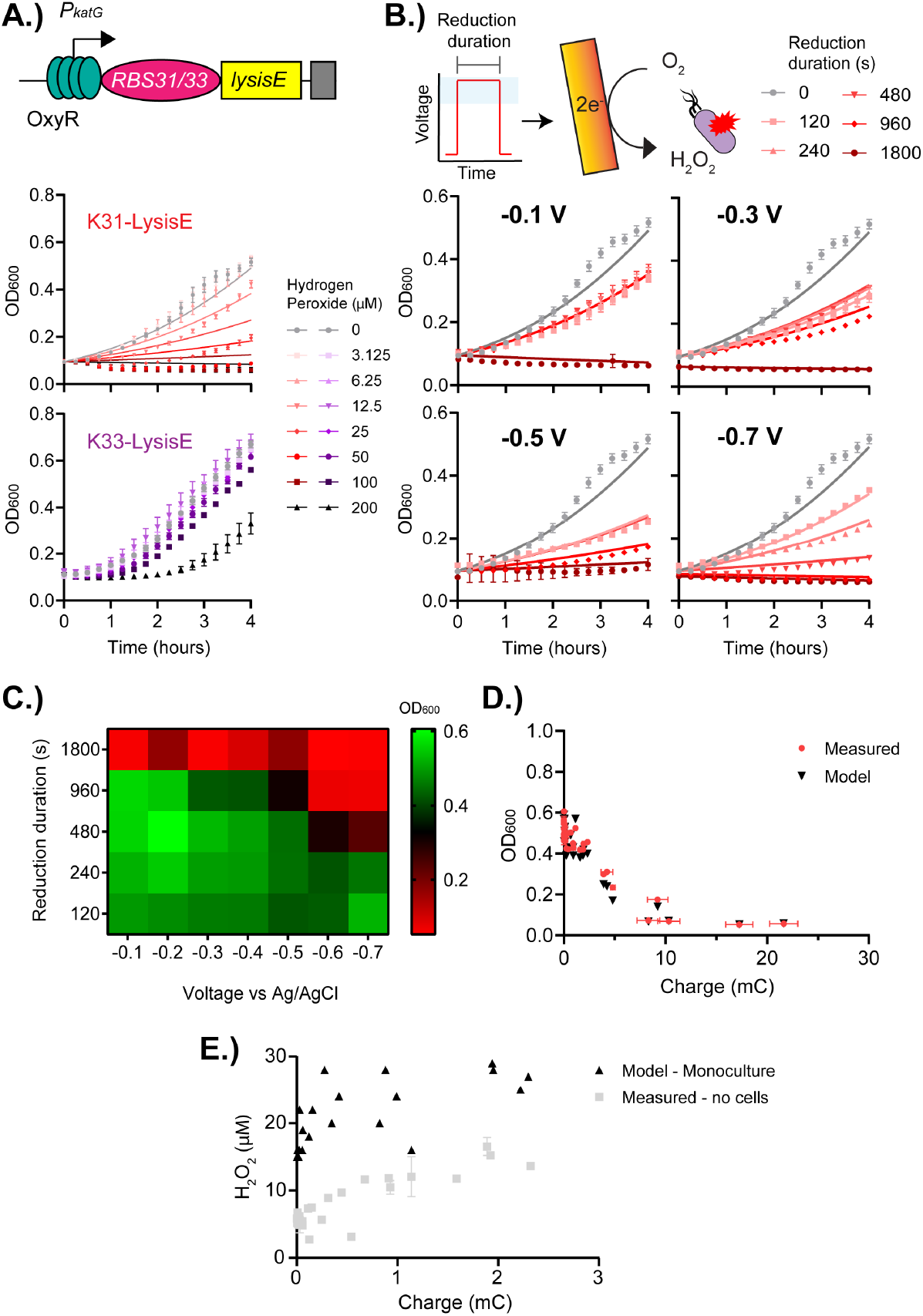
Direct electrogenetic induction of cell lysis within monocultures. **A.)** H_2_O_2_ sensitive gene circuits were constructed where LysisE expression is controlled by an OxyR activated *P*_*katG*_ promoter with either RBS31 or RBS33, designated K31-LysisE (top, red) and K33-LysisE (bottom, purple) respectively. Growth curves measuring optical density at 600 nm (OD_600_) over time for both strains are shown, where cells were induced (at time 0) with the indicated H_2_O_2_ concentration (between 0-200 μM) and cultured for 4 h. Solid lines for K31-LysisE curves correspond to model fits. **B.)** Electrogenetic induction links voltage and time inputs to H_2_O_2_ dependent induction of the K31-LysisE cells. Induction was performed by applying voltages between -0.1 V to -0.7 V and holding for 0 to 1800 s (see Figure S3 for complete dataset). Growth curves post induction are shown. Solid lines correspond to model fits. **C.)** Heatmap showing the final OD_600_ after 4 h of growth in B as functions of voltage inputs (horizontal-axis) and reduction durations (vertical-axis). Energy inputs increase towards the upper-right corner of the heatmap. **D.)** The final OD_600_ after 4 h growth in B plotted versus charge. The 1800 s, -0.1 through -0.3 V inputs were removed from this plot as outliers to the general trend. Experimental values, red circles; Model predictions, inverted black triangles. **E.)** Apparent initial H_2_O_2_ values from model predictions (black triangles) versus charge. Gray squares represent analogous, electrode produced H_2_O_2_ measurements (via colorimetric assay) without cells versus charge. All experiments were performed in triplicates.

Next (Figure 1B), we placed K31-LysisE cells in a previously developed electrogenetic induction chamber (VanArsdale et al., 2022) to determine if localized, time-dependent generation of H_2_O_2_ could induce a reduction in population density. These chambers catalytically produce H_2_O_2_ at different rates (measured H_2_O_2_ results are shown in Figure S2). We found at all applied voltages and for all durations, the OD_600_ of the affected cells was decreased. Most notably, the greatest decreases were observed for the -0.5 and -0.7 V cases where H_2_O_2_ generation is most efficient (Terrell et al., 2021). At these voltages and at durations above 480 s, there was the most charge (integrated current responses from the reduction duration and voltage inputs) and consequently the most H_2_O_2_ emitted from the electrode (Terrell et al., 2021). We extended the mathematical model for electrode actuation and results are again overlayed with the OD_600_ data. In Figure 1C, we mapped the OD_600_ data at 4 h showing the response profile to voltage and reduction-time respectively (raw data with more voltages are in Figure S3). Finally, in Figure 1D, we overlayed all data as a function of charge. Note that a given charge can be obtained given one voltage and different durations, as well as one duration and different voltages. Results show nearly linear reduction in final OD_600_ with applied charge until ∼10 mC wherein additional charge had no bearing on cell lysis, as all cells were dead.

Noting that the electrogenetically actuated cell death (Figure 1B) seemingly occurred at lower H_2_O_2_ levels (Figure S2B) than in experiments where known levels of H_2_O_2_ were added (Figure 1A), we decided to “fit’ our mathematical model to the data by creating an “apparent” H_2_O_2_ (Figure 1E) or an estimated H_2_O_2_ that when used as an initial condition, best fit our data. In nearly all cases, the “apparent” H_2_O_2_ was greater than what was observed even without cells (Figure S2B). This suggests that the extent of lysis was in fact greater than we would have expected based on generated H_2_O_2_. Interestingly, this was consistent with our previous work wherein H_2_O_2_ responsive cells, affixed to a gold electrode, needed a significantly lower concentration of H_2_O_2_ to activate genetic responses than that expected to be generated in the chamber volume at the same charge (Terrell et al., 2021). Moreover, our previous work also showed that electrogenetically-stimulated cells appeared to express more GFP than when provided an equivalent bolus of H_2_O_2_ to the medium (Terrell et al., 2021; VanArsdale et al., 2022). In all cases, our data reinforce the notion that cells in a mixed system transiently come in contact with the electrode, experience a higher H_2_O_2_ concentration than in the bulk solution, and respond more quickly and to a greater extent than predicted based on the bulk H_2_O_2_ level (VanArsdale et al., 2022). Here, the net effect was equivalent to a ∼2 to 4-fold increase in H_2_O_2_, suggesting a similar enhancement in signal transmission. We note also that the presence of activity metabolizing cells acts as a H_2_O_2_ sink, which would tend to attenuate the input signal and suggest that the apparent H_2_O_2_ is even higher than experimentally predicted.

Overall, these results indicate that cell lysis was efficiently induced when a reducing voltage was directly applied to OxyR responding cells. Moreover, our data reveals a more rapid and thorough response than the equivalent addition of H_2_O_2_ to the bulk solution.

### A transmitter-receiver model further reduces energy requirements and extends design options

Next, we built a “transmitter-receiver” system for signal propagation where one cell population transforms the H_2_O_2_ signal to QS autoinducer AI-1. In turn, AI-1 activates *lysisE* expression in another cell population and when sufficient LysisE has accumulated, cell lyse. Here, the “transmitter” cells contain an OxyR-activated gene circuit which regulates the expression of *lasI*, the gene encoding the 3-oxo-C12-homoserine lactone (AI-1) synthase (Mukherjee & Bassler, 2019). AI-1 produced via LasI is transported through the medium to “receiver” cells engineered to express *lysisE* via activation of *P*_*lux*_ by AI-1 bound transcription factor LasR (Tekel et al., 2019). Given the relative stability of AI-1 and LasR activation, the resultant communication therefore links the response of the receiver population to the proportion of the transmitter cells activated. This is analogous to cooperative thresholding in multicellular signaling, such that the response only proceeds when a sufficient number of cells are activated (Suderman et al., 2017). We hypothesized that signal relay would therefore result in a near “ON-OFF” response above a given H_2_O_2_ threshold. Consequently, we also expected this relay to lower the charge input requirement to cause lysis in the electrode system, as the information transmitted to receiver cells is energized by metabolic QS activity rather than an electrode.

In Figure 2A, we placed *lysisE* under the expression of the *P*_*lux*_ promoter alongside a strong constitutive expression cassette for LasR. We found that this “receiver” construct caused significant cell lysis between 0.78 to 3.13 nM AI-1, and a complete and sustained collapse in OD_600_ occurred when AI-1 was ∼6.25 nM. At AI-1 concentrations exceeding 6.25 nM, lysis occurred ∼30 min following AI-1 addition, suggesting LysisE accumulation was required for lysis. We named this construct LasR(WT)*-*LysisE, as it uses the wild-type LasR sequence. Based on a previously published model (Claussen et al., 2013; Welch et al., 2020), we extended our mathematical model to include LasR induction of *P*_*lux*_ and production of LysisE, accounting for dimerization of LasR and subsequent AI-1 binding for actuation. We parameterized the model using literature data and minimized the difference between our models’ predictions and experimental data (Supplemental methods). Our simulations (solid lines, Figure 2A) show that the model reasonably captured the AI-1 dependence of lysis, although lysis for cases above 6.25 nM appeared to lead the model simulations.

**Figure 2.**
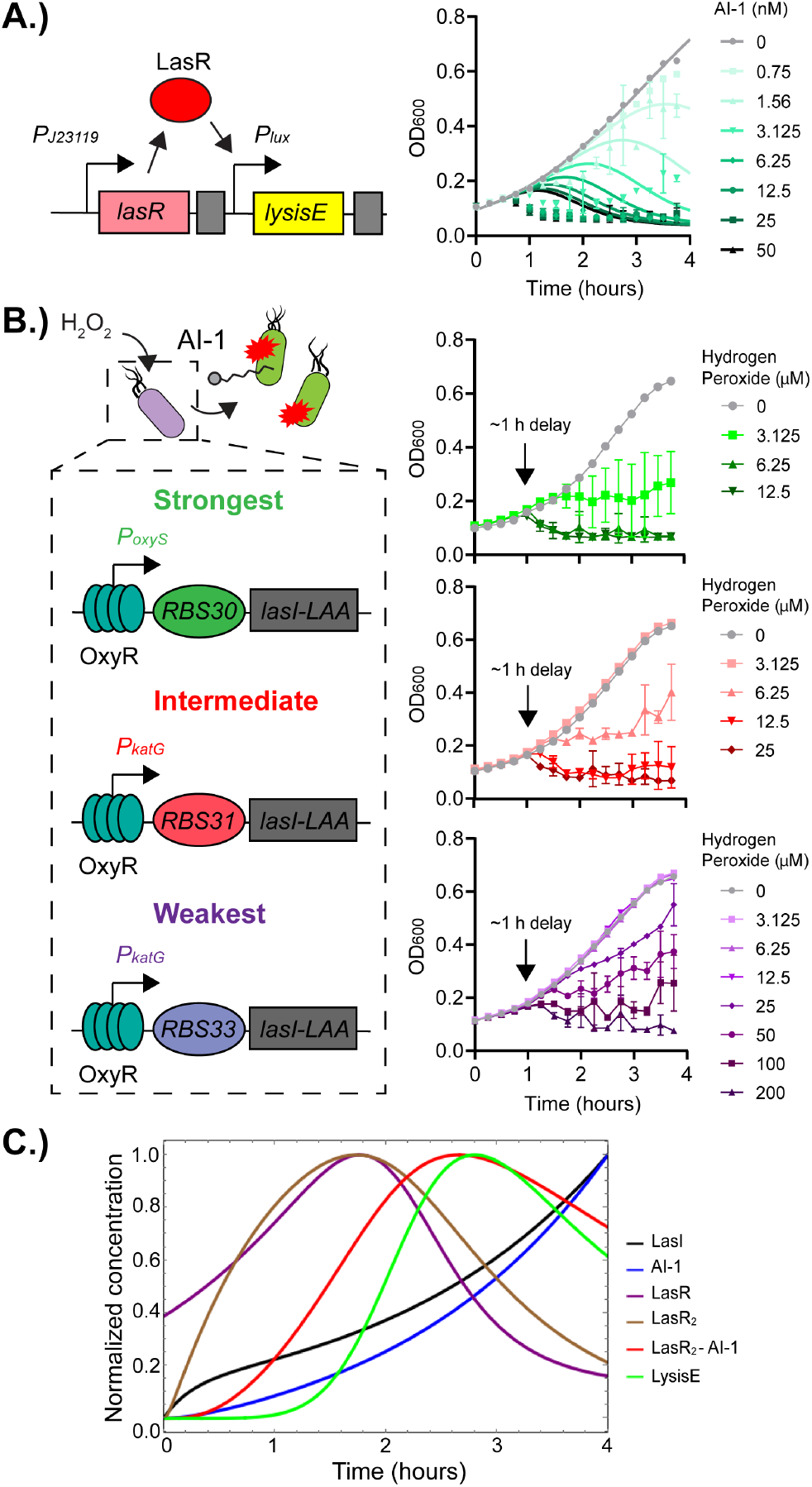
Designing a transmitter-receiver coculture for lysis of receiver populations. **A.)** Quorum sensing induced cell lysis was programmed into receiver cells by constitutively expressing LasR from the *P*_*J23119*_ promoter, which in turn activates *lysisE* expression from the *P*_*lux*_ promoter in response to AI-1. Growth curves measuring optical density at 600 nm (OD_600_) over time for receiver cells (LasR(WT)-LysisE, left) are shown, where cells were induced (at time 0) with the indicated concentrations of AI-1 (between 0-50 nM) and cultured for 4 h. Solid teal shaded lines correspond to model fits. **B.)** Growth curves for different transmitter-receiver cocultures are shown, where cells were induced (at time 0) with the indicated H_2_O_2_ concentration (between 0-200 μM) and cultured for 4 h. Cocultures initially contained LasR(WT)-LysisE receiver cells and 1% transmitter cells, either OxyR-LasI-LAA (“Strongest”, green, top), K31-LasI-LAA (“Intermediate”, red, middle), or K33-LasI-LAA (“Weakest”, purple, bottom) that are responsive to different H_2_O_2_ levels. All experiments were performed in triplicates. **C.)** Normalized concentrations of each component across the 4 h lysis window according to model fits for the OxyR-LasI-LAA transmitter system. LasR_2_ is LasR dimer. LasR_2_-AI-1 is AI-1 bound LasR dimer.

Next, we designed several “transmitter” constructs to link H_2_O_2_ to AI-1 generation. We expected this relay could be modulated through LasI production. To test this, we generated K31 and K33 transmitter constructs based upon the design in Figure 1A and a pre-existing OxyR-LasI construct with an SsrA degradation tag LAA on the C-terminus of LasI to reduce its lifetime; we have used this pre-existing construct in previous transmitter-receiver configurations (Stephens et al., 2021; VanArsdale et al., 2022) (Figure 2B, scheme). Both K31-LasI-LAA and K33-LasI-LAA variants were expected to produce less LasI than the *P*_*oxyS*_ construct. Each of the transmitter populations was then mixed with receiver cells at a total OD_600_ of 0.1 with the transmitter occupying 1% of the initial OD_600_. In preliminary experiments, we found this starting ratio prevented lysis of the receiver population, presumably due to minimal basal levels of AI-1 synthesis from the transmitters. As shown in Figure 2B, we found that, in agreement with prior experiments, the *P*_*katG*_ promoters had shifted the lysis dose-response by requiring greater concentrations of H_2_O_2_ to elicit similar decreases in OD_600_. That is, the weaker promoters presumably resulted in less AI-1 signal. The original OxyR-LasI-LAA transmitter construct caused a collapse in the coculture optical density when induced with 6.25 μM H_2_O_2_ (Figure 2B, green). By comparison, K31-LasI-LAA and K33-LasI-LAA caused complete lysis when induced with 25 μM and 200 μM H_2_O_2_, respectively (Figure 2B, red and purple). These concentration ranges mirror those seen in similar peroxide-dependent gene circuits published previously (Rubens et al., 2016) and were corroborated by fluorescent measurements using an equivalent mVenus AI-1 reporter circuit (Figure S4A). Compared to the monoculture communication model in Figure 1, the K31-LasI-LAA transmitter construct produced comparable OD_600_ attenuation at ∼4x less H_2_O_2_ (i.e., 12.5 μM versus 50 μM). Similarly, the comparable K33-LasI-LAA transmitter construct completely lysed the coculture above 200 μM, while the K33-LysisE construct in the monoculture system could not at any concentration. These results confirmed that the AI-1 production in a message propagation scheme amplified the H_2_O_2_ induction beyond the immediately activated cells. This is especially noteworthy, in that the proportion of the transmitters was low enough that outgrowth of the transmitters could not be detected within the experimental timeframe.

We subsequently examined model parameters for sensitivity and for maximizing agreement. The values that had the most corrective effect were first the constant for binding of H_2_O_2_ to the *P*_*katG*_ promoter via OxyR (k_1_) followed by the rate coefficient for production of AI-1 (k_fAI1_). The k_1_ values for K31-LasI-LAA and K33-LasI-LAA were 50% and 1% of that for OxyR-LasI-LAA, respectively. The k_fAI1_ values for K31-LasI-LAA and K33-LasI-LAA were analogously 75% and 66% of that for OxyR-LasI-LAA. These results corroborate our hypothesis that the changes to the promoter and RBS design decreased the expression of LasI (Figure S4B).

Interestingly, in all cases of the transmitter-receiver model, there was a noticeable ∼1 h delay from addition of H_2_O_2_ to lysis (arrows in Figure 2B), demonstrating the relay between each cell takes time to activate the different components of the system, reach the appropriate AI-1 threshold, and produce sufficient LysisE. This conclusion is supported by the normalized concentration dynamics in our mathematical model (Figure 2C), where the AI-1 bound LasR dimer complex (LasR_2_-AI-1) preceded the rise in LysisE. In this figure (modeled for the OxyR-LasI-LAA transmitter system), the expected temporal switching was observed with LasI accumulation preceding AI-1, followed by LasR and LasR dimer (denoted LasR_2_). Finally, LasR_2_-AI-1 preceded LysisE production, which began at ∼1 h post addition of H_2_O_2_. The model enables rationalization of the principal differences between the monoculture system and the transmitter receiver system: (i) the time delay associated with AI-1 synthesis and transport and (ii) the interaction of the LasR dimer with AI-1 (instead of direct H_2_O_2_ actuated *lysisE* expression). In both cases, LysisE must accumulate, presumably identically to initiate lysis.

We next tested whether similar dynamics would be observed when using an electronic signal to generate H_2_O_2_. As schematically depicted in Figure 3A, the OxyR-LasI-LAA transmitter cells present in the stirred electrochemical cell chamber come into proximity of the electrode and are thus exposed to higher H_2_O_2_ concentrations than the bulk. In turn, these cells (also present at 1% of the total OD_600_) should respond exactly as in Figure 2 following their detection of H_2_O_2_, wherein they generate AI-1 and stimulate lysis in receiver cells. Interestingly, the dynamic responses of the population exhibited a shorter delay in lysis when compared to results with simple bulk addition of H_2_O_2_ (Figure 2). The complete set of OD_600_ results from varied input voltages and reduction times are found in Figure S5A. We suspect that the shorter time lags stemming from electronic induction are again aligned with transient exposure of transmitter cells to high H_2_O_2_ levels and subsequent induction of LysisE. That is, if even a few cells are exposed early to high levels of H_2_O_2_, a rapid signal transmission to the receiver population can follow. Figure 3B shows the response profile after 4 h of growth. The most dramatic attenuation was at -0.7 V and reduction durations exceeding 960 s. We note that even at the shortest reduction duration (i.e., 120 s), we observed fairly complete reduction in OD_600_. The final OD_600_ values after 4 h are also depicted in Figure 3C as a function of applied charge. Here, only 2.5 mC were needed to elicit nearly complete lysis. This is only 25% of that needed for the monoculture model, albeit with a less sensitive H_2_O_2_ circuit (Figure 1). This result suggests QS signal relay dramatically reduced the originating energy input needed to establish lysis in the receiver population.

**Figure 3.**
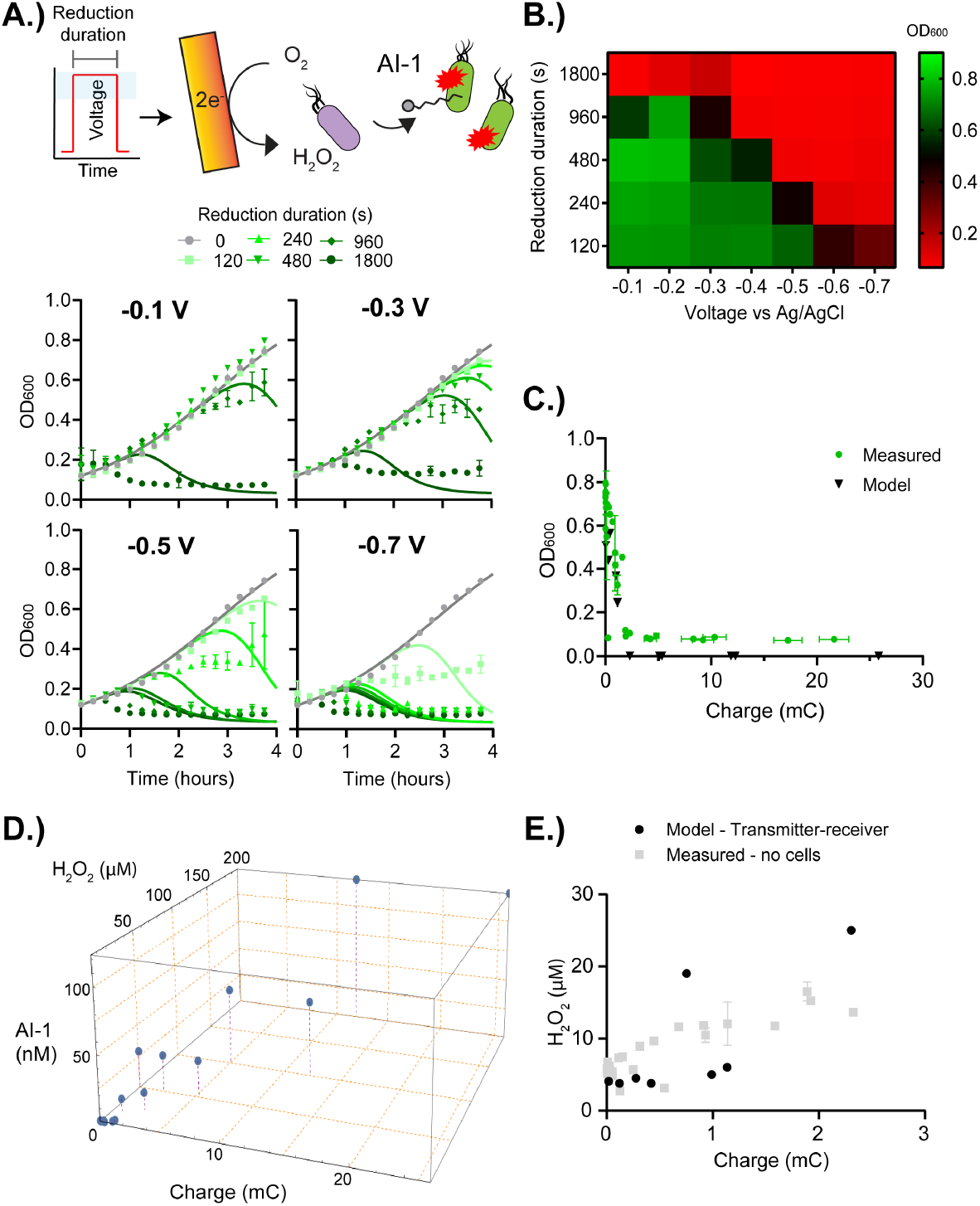
Electrogenetic induction of OxyR-LasI-LAA transmitter-receiver cocultures lyse the receiver population. **A.)** Electrogenetic induction links voltage and time inputs to H_2_O_2_ dependent induction of the OxyR gene circuit within the transmitter cells, which in turn produces AI-1 to lyse the receiver population. Induction of transmitter-receiver cocultures initially containing 1% transmitter cells was performed by applying voltages between -0.1 V to -0.7 V and holding for 0 to 1800 seconds (see Figure S5 for complete dataset). Growth curves post induction are shown. Solid lines correspond to model fits. **B.)** Heatmap showing the final OD_600_ after 4 h of growth in A as functions of voltage inputs (horizontal-axis) and reduction durations (vertical-axis). Energy inputs increase towards the upper-right corner of the heatmap. **C.)** The final OD_600_ after 4 h growth in A plotted versus charge. The 1800 s, -0.1 through -0.3 V inputs were removed from this plot as outliers to the general trend. Experimental values, green circles; Model predictions, inverted black triangles. **D.)** The model prediction of the relationship between charge, produced AI-1, and the “apparent” initial H_2_O_2_ concentration that induces the transmitter population. **E.)** Apparent initial H_2_O_2_ values from model predictions (black circles) versus charge. Gray squares represent analogous, electrode produced H_2_O_2_ measurements (via colorimetric assay) without cells versus charge. All experiments were performed in triplicates.

To further our analysis, we used our mathematical model to back calculate both “apparent” initial H_2_O_2_ concentrations (experienced by transmitter cells, assuming a bolus of H_2_O_2_ addition) and AI-1 concentrations that best fit the OD_600_ data (Figure 3A) and plotted these values versus the input charge (Figure 3D, E, Figure S5B, C). At very low charge (Figure 3E), we found that the apparent amount of H_2_O_2_ generated exceeded 6.25 μM, suggesting that the induction of AI-1 synthesis was more than sufficient to activate the *lysisE* gene in the receiver cells at nearly all voltages and durations. It is interesting that for the monoculture model (Figure 1E), the apparent H_2_O_2_ concentration was higher than expected (owing to gained efficiency of distributed exposure to H_2_O_2_) while the transmitter-receiver model under-predicted the concentration (likely owing to a signal amplification converting the original electronic input to QS-based molecular signaling). This latter amplification result is likely because the QS signal has substantially different chemical properties than H_2_O_2_ and consequently appears to be more effective at signal transmission. For example, with AI-1 transmission, there was a reduced requirement for H_2_O_2_ so that the transmitters were less impacted by exposure to H_2_O_2_ that, in turn, negatively affects cell physiology (Fu et al., 2015). Moreover, H_2_O_2_ is rapidly degraded, necessitating its continual production. AI-1, by contrast, is continually produced by LasI (long after the initial transmission) and because it is not actively degraded in our system, it persists, enabling accumulation and continued message propagation.

### Genetic circuits add diversity to design, enabling optimization of signal propagation

As noted above, another interesting difference between the monoculture and transmitter-receiver systems was that the transmitter-receiver model typically exhibited a ∼1/2 to 1 h time delay prior to lysis. Our model suggests this is due to LasR_2_-AI-1 dimer accumulation, resulting in delayed LysisE expression. We decided to experimentally test this conclusion by creating genetic constructs that decrease the affinity for AI-1 binding that, in turn, would negatively impact LasR-directed expression. Bassler and coworkers had previously shown that S129 LasR mutations altered the EC_50_ value for dimer association with AI-1, in their work delaying subsequent pyocyanin synthase expression in *Pseudomonas aeruginosa* (McCready et al., 2019). We hypothesized this same approach could be used to delay the onset of lysis by elevating the concentration of AI-1 required for LysisE expression and thereby delay its accumulation and function. We selected mutations S129W, S129F, S129T, and S129M, of the Bassler work which were reported to have EC_50_ values of 77 nM, 177 nM, 870 nM, and 6600 nM for AI-1, respectively (McCready et al., 2019). We built on their results by inserting *lysisE* in expression vectors with identical mutations in LasR enabling varied LysisE synthesis in AI-1 responsive receiver cells. In Figure S6, we found that cells carrying S129W lysed above 100 nM of supplemented AI-1, and both the S129F and S129T mutations lysed above 1000 nM. These values corresponded nicely to the Bassler work (McCready et al., 2019). Note, we did not observe lysis for the S129M variant at the AI-1 levels tested.

Next, we evaluated the effect of LasR mutation on AI-1 mediated expression by replacing *lysisE* with the gene for expression marker, CFP. Shown in Figure S7A are fluorescence results from experiments in which H_2_O_2_ was added to the transmitter-receiver coculture wherein the transmitter population was OxyR-LasI-LAA and the receiver population was replaced by the varied LasR mutants making CFP. These experiments show how the H_2_O_2_-stimulated synthesis of AI-1 was perceived by the receiver population. In all cases, the S129 mutations clearly attenuated CFP expression at all levels of H_2_O_2_ and the degree of attenuation corresponded with increased EC_50_ of the LasR mutant. These experiments indicate that upon H_2_O_2_ addition, coculture experiments will exhibit a delayed lysis that is roughly dependent on the amount of time needed for the requisite AI-1 LasR binding to occur, and naturally, this is attributed to transmitter AI-1 synthesis, AI-1 transport, and its accumulation and binding within the receiver cells.

In Figure 4A, we tested the complete transmitter-receiver model wherein the receiver cells generated LysisE as directed by the mutant LasR activators. The results were as anticipated. At time = 0, we added H_2_O_2_ to transmitter-receiver cocultures where the transmitter cells were the OxyR-LasI-LAA cells and the receivers contained either the LasR(S129W)-, LasR(S129F)-, or LasR(S129T)-LysisE variants. For comparison, we added shaded regions onto the plots to highlight the points at which the OD_600_ first appeared to deviate from the uninduced controls. Recall that LasR(WT)-LysisE was ∼1 h post induction (Figure 2). By comparison to the WT circuit, LasR(S129W)-LysisE seemingly delayed lysis by an additional ∼30 minutes and LasR(S129F)-, and LasR(S129T)-LysisE each delayed lysis by an additional ∼1 h (Figure 4A). These delay times are supported by “time of onset” values (see Methods) that we calculated based on data interpolation (Figure 4C). Generally, we find that higher concentrations of H_2_O_2_ were needed to achieve lysis in the LasR variants. Notably, 6.25 μM H_2_O_2_ was needed to halt growth in LasR(S129W)-LysisE, while 12.5 μM and 50 μM were needed to suspend OD_600_ accumulation in LasR(S129F)- and LasR(S129T)-LysisE, respectively. Also, in both LasR(S129F)- and LasR(S129T)-LysisE, even at 200 μM H_2_O_2_, we found no rapid decrease to the preinduction level. Instead, we found an array of “intermediate” responses between completely unperturbed and various extents of lysis as a function of the H_2_O_2_ added.

**Figure 4.**
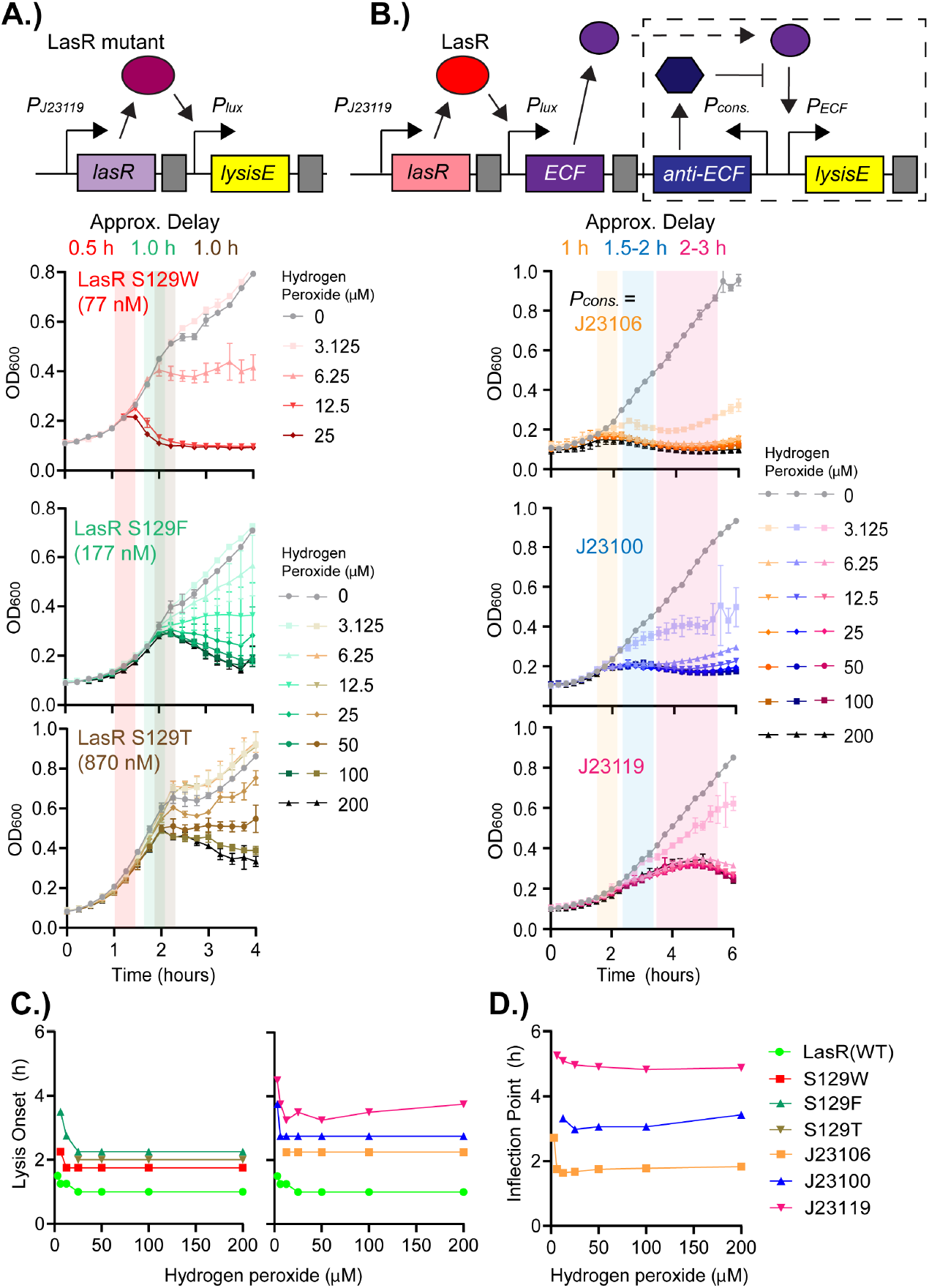
Programming a time delay in lysis response. **A.)** A time delay was introduced into receiver constructs by mutating the S129 codon of the LasR transcriptional activator, which lowers its affinity for AI-1 such that it does not become active until higher concentrations of AI-1 have accumulated. Growth curves for transmitter-receiver cocultures are shown, where cells were induced (at time 0) with the indicated H_2_O_2_ concentration (between 0-200 μM) and cultured for 4 h. Cocultures initially contained the indicated LasR mutant receiver cells (AI-1 EC_50_ values in parentheses) and 1% OxyR-LasI-LAA transmitter cells. The shaded boxes indicate the approximate time of lysis of each variant. The approximate delays relative to the LasR(WT) variant, which lyses at ∼1 h post induction, are as follows: ∼0.5 h for LasR(S129W) (top, red); ∼1 h for LasR(S129F) (middle, green) and LasR(S129T) (bottom, brown). **B.)** A transcriptional time delay was introduced into receiver constructs by creating an ECF/anti-ECF competitive antagonist relationship in which ECF, expressed in response to AI-1, must overcome an antagonistic threshold of anti-ECF to sufficiently express LysisE and induce lysis. The threshold of anti-ECF was altered using constitutive Anderson promoter variants (in order of increasing strength): J23106, J23100, and J23119. Growth curves of transmitter-receiver cocultures were conducted as in A and the shaded boxes indicate the approximate time of lysis. The approximate delays relative to the LasR(WT) variant are as follows: ∼0.5 h for J23100 (top, orange), ∼1-1.5 h for J23100 (middle, blue), and ∼2-3 h for J23119 (bottom, pink). **C.)** The Lysis Onset times for each LasR variant (left) and ECF/anti-ECF variant (right) were plotted versus induction concentration. A Lysis Onset time was calculated as the time at which the difference between the null control (average grey lines in each plot) and the induced tests (average colored lines) exceeded a threshold of 0.1. **D.)** The Inflection Points for each ECF/anti-ECF system were plotted versus induction concentration. The Inflection Point was calculated as the time at which the derivative of the average response became negative, thus indicating the population was decreasing. All experiments were performed in triplicates.

In sum, these results indicated that mutating LasR could, by enabling distinct alterations in AI-1 binding, change population dynamics in a predictive manner by delaying the onset of lysis. Interestingly, we found the delays introduced were somewhat modest (∼1 additional hour), given the necessary rise in AI-1 might have exceeded 10-fold with 5-fold differences in EC_50_ values. Moreover, LasR(S129F)-LysisE and LasR(S129T)-LysisE system behaviors were quite similar. Perhaps either (i) the concentration rise of AI-1 between 100-1000 nM occurs rapidly or (ii) partial *P*_*lux*_ activation at sub-EC_50_ values is sufficient to cause a deviation in population trajectory.

While altering just one component, a transcriptional regulator, in the overall receiver circuit did indeed enable a predictable shift in consortia composition, we further hypothesized that additional, more complex genetic circuits could achieve more dramatic shifts, both in time and amplitude. In Figure 4B, we introduced a competitive interaction between a heterologous transcriptional activator (extracytosolic factor 20, ECF20 (Rhodius et al., 2013)) and an antagonist (anti-ECF20) that had been previously shown in both *E. coli* (Chen & Arkin, 2012) and a cell-free system (Shin & Noireaux, 2012) to achieve delayed actuation. This addition insulates LysisE expression until a “capacitive threshold” can be overcome by a signal-relay carrier. Here, we placed the expression of ECF20 under the control of the *P*_*lux*_ promoter, and then placed the expression of *lysisE* under the *P*_*ECF*_ promoter. We then altered the expression level of anti-ECF20 using different strength constitutive Anderson promoters (iGEM Registry of Biological Parts, http://parts.igem.org/Promoters/Catalog/Anderson). In all figures, each promoter tested is listed top-to-bottom in order of increasing normalized strength such that J23119 produces the most anti-ECF and J23106 produces the least, correspondingly offering the most and least repression of ECF activation.

Interestingly, we found that newly constructed delay circuits employing J23106, J23100, and J23119 delayed lysis in the transmitter-receiver model by an additional 30 minutes, 1-1.5 hours, and 2-3 hours, respectively (Figure 4B). As anticipated, the delay corresponded with anti-ECF promoter strength. Based on data interpolation, we find that the onset of lysis was extended and here, also the point of inflection time (Figure 4C, D). Note, these two times were more distinct in these cases compared to the LasR mutant systems, resulting in the wider shaded regions in Figure 4B. This was most evident in the J23119 variant, which inhibited growth starting at 2 h, but did not decrease OD_600_ until about 5 h. The calculated time of onset and inflection times indicate the extent to which altered genetic designs elongate and attenuate signal propagation, enabling the capacity of a more fine-tuned outcome. Thus, we suggest that circuit activation, which is regulated by the LasR(WT), “tunes” the concentration sensitivity for initiating ECF20 transcription, while the promoter strength of the anti-ECF20 “tunes” the eventual accumulation of LysisE.

These results are supported by analogous fluorescence experiments where we replaced *lysisE* with the gene for CFP in order to fully characterize the *P*_*ECF*_ expression without interference from lysis. In each case, the rise in fluorescence was both reduced and delayed corresponding to the anti-ECF promoter strength (Figure S8). These trends were supported by Fritz and coworkers (Pinto et al., 2018). Interestingly, the relative fluorescence units (RFUs) were significantly greater than those of the LasR variants in Figure S7, indicating that the “capacitive threshold” design of the *P*_*ECF*_ promoter and ECF20 together are more efficient at initiating expression than the native *P*_*lux*_ system. We note at the highest levels of H_2_O_2_, there is a decrease in fluorescence, perhaps due to high CFP overexpression inducing burden on metabolism. An alternative contributing factor might have been hydrogen peroxide toxicity at levels above 50 μM.

Finally, we linked both the LasR variant and ECF/anti-ECF systems to the electrogenetic induction system to determine how QS propagation of the signal would proceed in these modified contexts (Figure 5A). To do this, we repeated identical set of experiments from Figure 4 but with electrode generated H_2_O_2_ and over the full range of applied voltage and duration. Complete results, depicted in Figures S9 and S10A, are summarized in Figure 5B-C as a function of charge. In the top subpanels of Figure 5, we depict results of LasR(S129W) and (LasR)S129F (Figure 5B) and J23106, J23100, and J23119 (Figure 5C) receiver systems. First, by comparison to analogous plots in Figures 1D and 3C, the data here are more scattered. Perhaps this reflects the greater number of trajectories possible with distributed signal transmission at the electrode and the altered genetic circuits that extend signal propagation. For example, these results are quite different than when H_2_O_2_ was applied in a bolus, indicating the charge-to-QS relay significantly impacted actuation of these circuits. We note that for the LasR(S129F) case, there was no significant lysis irrespective of charge (Figure S9). For the LasR(S129W) mutant, we found >15 mC charge was needed to obtain significant cell lysis compared to uninduced tests. Thus, in these cases, we found more charge was needed to effect a similar change as the H_2_O_2_ addition experiments. This was as anticipated and interestingly, the changes in delay time relative to the H_2_O_2_ addition experiments were not very significant. Instead, the sensitivity to charge was more substantially affected. In the case of the ECF/anti-ECF delay circuits, we found results were more consistent with the experiments of added H_2_O_2_, with higher and more sustained OD_600_ for the circuits with longer delays. This was perhaps because the capacitive threshold results in a more all-or-nothing response relative to the LasR variants, which have a greater spread in sensitivity to transmitter activation compared to the WT.

**Figure 5.**
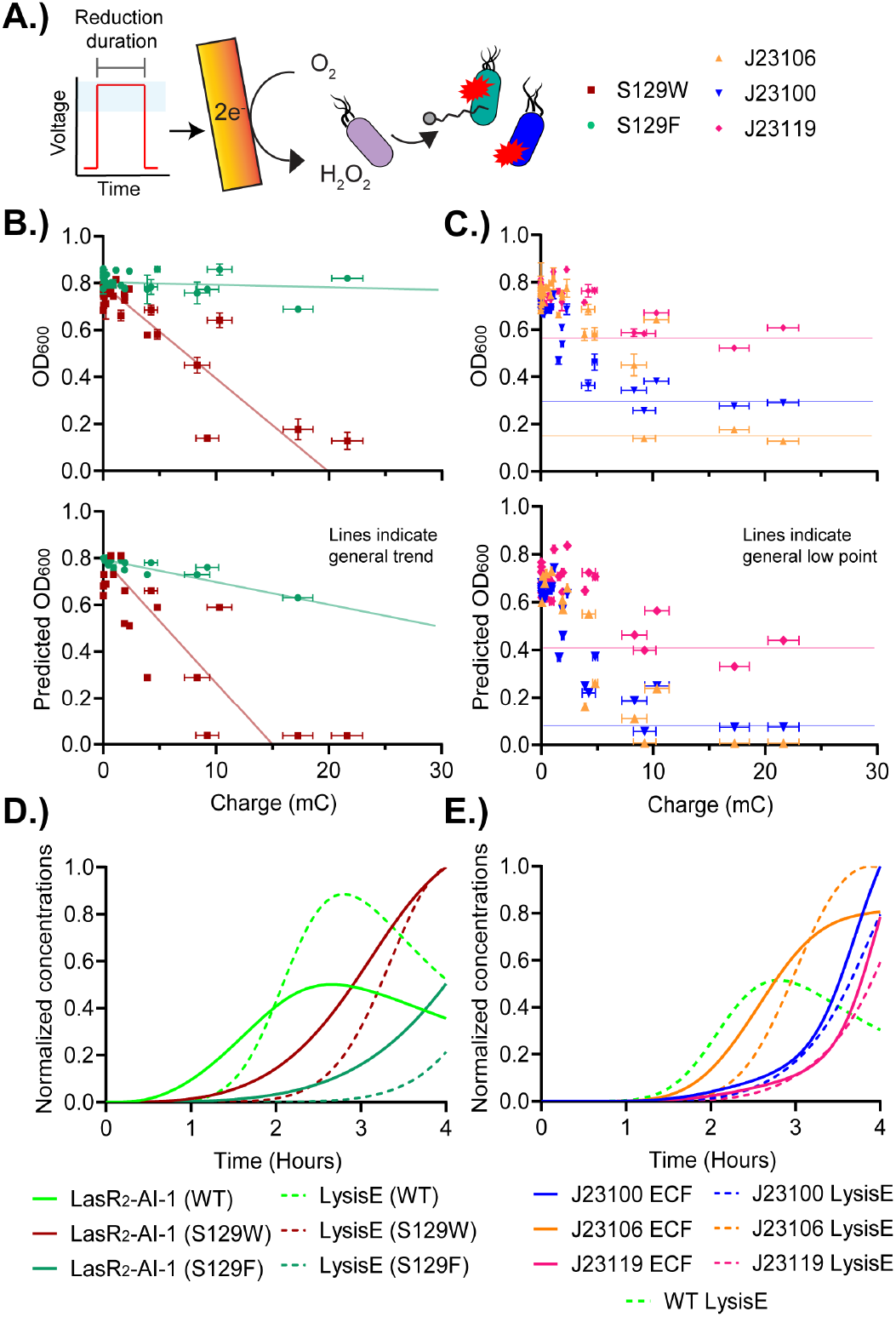
Electrogenetic activation of LasR mutant and ECF/anti-ECF delay systems. **A.)** Electrogenetic transmitter-receiver coculture induction was performed as described in Figure 3A using indicated LasR and ECF/anti-ECF variant receivers and OxyR-LasI-LAA transmitters. **B-C.)** The final OD_600_ after 4 h growth following electrogenetic induction plotted versus charge (see Figures S9 and S10 for growth curves and heat map). Coculture data with LasR receiver variants are shown in B, while data with ECF/anti-ECF receivers are presented in C. Experimental measurements are shown in the top panel, while model predictions are shown in the bottom panel. Lines added in B indicate the relative trajectory trendlines. Lines added in C highlight the relative low-points of each charge-cell density relationship at the final measurement point. **D-E.)** Normalized concentrations of each component across the 4 h lysis window according to model fits for the LasR mutants (D) and the ECF/anti-ECF delay systems (E). LasR_2_-AI-1 is AI-1 bound LasR dimer.

To further investigate this dynamic, we fit our model of these systems to match “apparent” H_2_O_2_ to the input charge for each system. Our model predictions (bottom subpanels of Figure 5B) of the relationship between final OD_600_ to charge confirmed that altering the LasR variant substantially increased the amount of charge needed to achieve a given lysis level. Models of the LasR(S129W) receiver system decreased in accordance with the data, both indicating general decrease in final OD_600_ with charge and the lowest OD_600_ data occurred in the vicinity of ∼20 mC. This is roughly a 10-fold increase relative to the WT system, again illustrating the need for additional power input when genetic delays were included. Also, the lower affinity LasR variant systems did not lyse, suggesting these receivers required greater charges than those tested. For the anti-ECF/ECF systems (Figure 5C), the model predictions and experiments were also in agreement. The OD_600_ versus charge relationships of the J23106 and J23100 constructs appeared similar until ∼5 mC, after which J23100 leveled off while J23106 continued to decrease with charge until ∼23 mC. Colored lines in Figure 5C show the final OD_600_ reached by each system, and these reflected the relative strength of the anti-ECF promoter.

We next show representative model results to help understand these system dynamics relative to the LasR(WT) transmitter-receiver system. In Figure 5D, as expected, we found that the expression of LysisE was delayed relative to the WT or mutant LasR_2_-AI-1 dimer complex. Moreover, the shift appeared to be about an hour in each LasR variant case. Because AI-1 accumulates at the same rate in all cases based upon the input signal (i.e., H_2_O_2_ or charge), the 1 h delay is compounded by the amount of time required for AI-1 to reach the appropriate threshold for each LasR variant. By comparison, in Figure 5E we see that the expression of ECF is closely tied to the onset of LysisE production. This partially occurs because the *P*_*ECF*_ promoter was observed to be substantially stronger than the *P*_*lux*_ promoter (Figures S7, S8). Therefore, the anti-ECF concentration, determined by the relative strength of the constitutive Anderson promoter, plays the predominant role in delaying transcription of *lysisE* because it prevents the accumulation of free ECF available. It is noteworthy, however, that ECF transcription is initiated immediately following formation of the LasR_2_-AI-1 complex as can be observed by comparing the weakly suppressed J23106 variant (orange line) to the accumulation of the WT *lysisE* (dashed green line). Finally, the model simulations well reflect the calculated time delays in that the delays for the LasR mutants were typically shorter than that for the ECF/anti-ECF constructs, and further that the delays for the latter followed anti-ECF promoter strengths. Together, our model and experimental results demonstrate how the confluence of growth dynamics (i.e., which impact availability of promoters and generation of activators like ECF) and threshold sensitivities to an inducer come together to determine the effectiveness of a signal-relay system.

## Discussion and Conclusions

Here we demonstrate that programmatic lysis of subpopulations within a microbial consortium through electrogenetic induction signals is dependent on how the signal is perceived and propagated through a network of both extracellular and intracellular interactions. We found that populations that were induced through direct perception of electrochemically produced H_2_O_2_ (the monoculture model) required long signal-durations to effectively activate the lysis switch within the majority of the population. By comparison, we found that transformation of the electrogenetic signal to a QS signal based on metabolic energy supply (the transmitter-receiver model) dramatically reduced both the time and input energy requirements needed to activate lysis in a majority of the population. This was done without altering the sensitivity of the H_2_O_2_-perceptive population to the incoming signal. These results are most evident by comparing the plots of 4-h OD_600_ data for each model versus charge. The applied charge is the closest equivalent to a true measure of energy input into the system. Our mathematical model reveals the shift away from H_2_O_2_-mediated gene activation to the interaction between AI-1 and LasR. This result is important to consider when designing electronic induction schemes in general. For example, in manufacturing systems, cells that are not induced within a batch culture lower the product yield per biomass and product yield per substrate, thereby necessitating some form of cell-cell communication or gratuitous induction for reestablishing high productivity.

Integrating QS with electrogenetics revealed two interesting dynamics relevant to consortia signal relay that we want to highlight. First, the relay of the electrode-produced H_2_O_2_ to AI-1 through the transmitter population increased the lysis outcome relative to the proportion of the number of cells that were activated. In the monoculture system, increasing the input charge proportionally decreased the overall cell growth rate. This occurred because both the input signal transmission ceases due to lysis and it is degraded by all cells (i.e., those carrying data and those not). By contrast, increasing the input charge in the transmitter-receiver system was quite effective in transmitting the lysis directive. We observed that when activated above a given threshold, an “all-or-nothing” collapse in cell density occurs despite only 1% of the culture containing a circuit sensitive to the electrode signal. This ultimately occurred because signal reception by low-density transmitters continually propagated the signal through AI-1, and these cells did not lyse during the process, ensuring that the signal could not be terminated. The AI-1 reception by the receiver population, in turn, amplified the total *lysisE* expression. Second, we find that altering the response of the receiver to AI-1 ultimately influenced the input costs to achieve comparable levels of lysis to the original WT transmitter-receiver system. This observation was tested using LasR variants wherein the amount of AI-1 required for receiver activation was altered causing changes in *lysisE* expression. By comparison, however, the addition of the ECF/anti-ECF capacitive elements ultimately suppressed the consequences of signal reception, such that the accumulation of LysisE did not occur until later in the growth curve. As a consequence, fewer receiver cells lysed, indicating that the suppression of the *lysisE* transcription rate affected the sensitivity of the overall system to input charge. This effect is reminiscent of the role rate processes can play in noise suppression (Lestas et al., 2010), in that the capacitive threshold is reducing the desired action of the receiving cell but ultimately not its ability to receive the incoming AI-1 signal. Formal information theory analyses could help tease out exactly how these elements function and produce further generalizable conclusions in terms of input/output relationships within signal propagation chains (Harper et al., 2018; Pierobon & Akyildiz, 2013). Overall, these signal relay features are important considerations when designing synthetic consortia reliant on transient environmental cues for function (e.g., probiotic plant growth, pathogen suppression, bioremediation, biocontainment).

Future applications of electrogenetic lysis systems will be useful for illuminating the physiology of microbial consortia. Isolating specific members from bacterial communities and reintroducing them with responsive components (Rubin et al., 2022) should identify various design constraints for designing altruists for probiotic intervention (Ikryannikova et al., 2019) or as a means to isolate persistors (Gollan et al., 2019). Specially designed microfluidic systems could also be used to model the interactions of consortia with redox signals. For example, in a simple set-up with artificial soil, an electrode could be viewed as a model root releasing H_2_O_2_. Using a combination of electronic input and QS-based synthetic biology should enable evaluation of engineered synthetic soil communities prior to plant-based experiments (Liu et al., 2019). In an exactly analogous manner, wound or infection abatement could be studied as microsystems where geometrically defined electrodes could be created to mimic the physiological system. Further, such an electrogenic/QS signal propagation system could prove beneficial for evaluating biofilm growth, especially when the microbes are naturally electroactive (Logan et al., 2019). Future systems could then take advantage of biofilms in detection and remediation, such as within local water supplies (Atkinson et al., 2022). Ultimately, we believe these tools will assist the development of living microbiome materials that participate in a greater internet of information exchange (Akyildiz et al., 2015).

## Author Contributions

Preparation of manuscript (EV, AN, GP, WB, MCY); Performed experiments (EV, MC, TH, C-YC, SW); Computational modeling (AN); Research oversight (GP, WB, AN, YJ, MCY)

## Acknowledgements

Financial support of this work was provided in part by the National Science Foundation (CBET#1932963, ECCS#1807604, CBET#1805274), the Defense Threat Reduction Agency (HDTRA1-19-1-0021), the LLNL Laboratory Directed Research and Development program (17-ERD-013 to MCY), and the U.S. Department of Energy (DOE), Office of Science, Office of Biological and Environmental Research, Lawrence Livermore National Laboratory SFA “From Sequence to Cell to Population: Secure and Robust Biosystems Design for Environmental Microorganisms.” This work was performed under the auspices of the U.S. DOE by Lawrence Livermore National Laboratory under Contract DE-AC52-07NA27344 (LLNL-JRNL-841967).

