## Supplemental Information for "Electrogenetic signaling and information propagation for controlling microbial consortia via programmed lysis"

### Supplemental Methods and Modeling

*Fluorescent measurements.*

Cells were induced as described above either with hydrogen peroxide or ORR electrogenetic induction, as well as through addition of AI-1. Cell growth post-induction and fluorescence measurements were conducted in TECAN Spark plate reader pre-warmed to 37°C with linear shaking in a humidity-controlled chamber. Fluorescent measurements of mVenus (Ex: 515 Em: 527) and sCFP(3A) (Ex: 433 Em: 474) were measured every 15 minutes for 16 hours.

*Cell Preparation for Scanning Electron Microscopy.*

The K31-*lysisE* cells were cultured in minimal medium and induced with 200 µM $H_2O_2$ for 1 h. Cells were then collected by centrifugation at 3,000 x g for 3 min, washed twice in water and resuspended in 25% ethanol. After 30 min at room temperature, the cells were centrifuged (3,000 x g for 3 min) and resuspended in 50% ethanol. Two more solvent exchanges in 75% and 100% ethanol were performed using the same centrifugation and incubation steps. Cells suspended in 100% ethanol were drop-cast onto foil, dried, and subsequently gold sputtered (5 nm layer), then imaged on the Quanta 250 FEG (FEI company; Hillsboro, OR) at a beam voltage of 30 kV.

*Model Methods and Equations.*

*Monoculture system:*

We simulated all cell growth using the Verhulst model.

$$\frac{dT(t)}{dt} = T(t)\left(k_T\left(1 - \frac{T(t)}{T_\infty}\right) - k_{dt}\right) \qquad (1)$$

Where

$T(t)$ = concentration of transmitter cells at time $t$

$k_T$= carrying capacity coefficient for $T$

$T_a$ = carrying capacity of $T$ in the medium

$k_{dt}$ = natural death rate coefficient for both transmitter and receiver cells

To simulate "monoculture" electrogenetic cell lysis with K31-LysisE (Figure 1B), we treated $H_2O_2$ as a non-competitive growth inhibitor. To match the observed behavior for small concentrations of $H_2O_2$, we added a fast sink reaction whereby up to 6.25 µM of $H_2O_2$ is removed from the system almost immediately after simulation begins. We used the results for direct $H_2O_2$ induced expression (Figure 1A) to calculate an $H_2O_2$ inhibition constant ($K_{i,H2O2}$) for K31-LysisE.

$$\frac{dT(t)}{dt} = T(t)\left(k_T\left(1 - \frac{T(t)}{T_\infty}\right)\left(\frac{1}{\left(1+\frac{H_2O_2(t)}{K_{i,H_2O_2}}\right)}\right) - k_{dtH_2O_2}H_2O_2(t)\right) \qquad (2)$$

From Figure 1A, we observe that high concentrations of $H_2O_2$ inhibit growth, indicating toxicity. Thus, we added the last term in eq. (2) to account for the toxicity of $H_2O_2$.

To calculate the inhibition constant, as well as other model-based constant predictions, we

1. altered the parameter in the model,
2. calculated the sum square of error (SSE) between model prediction and experimental measurements, and
3. continued altering the parameter until a minimum SSE was reached or the SSE curve practically flattened.

We used the results from eq. (2) to predict $H_2O_2$ concentrations that would result in similar growth inhibition as those induced by electrogenetic methods. For simulations in Figure 1B, for each of set of measurements, we simulated eq. (2) while varying the values of $H_2O_2$ to ensure minimal SSE. The predicted "apparent $H_2O_2$" values from these simulations are shown in Figures 1E and S3B.

For the remainder of the simulations we converted the $OD_{600}$ values to nM values using the conversion factor 1 $OD_{600}$=8x$10^8$ cells/ml= 0.0013 nM from the Agilent biocalculator (www.agilent.com/store/biocalculators/calcODBacterial.jsp). The growth rate coefficients for each simulation were calculated by minimizing SSE for the system without any induction of lysing.

*Transmitter-receiver systems:*

For the LasR(WT)-LysisE system, the receiver cells growth was also simulated using the Verhulst model.

$$\frac{dR(t)}{dt} = R(t)\left(k_R\left(1-\frac{R(t)}{R_\infty}\right) - k_{dt} - k_{lys}Lys(t)\right) \qquad (3)$$

Where

*R(t)* = concentration of receiver cells at time *t*

$k_R$= carrying capacity coefficient for *R*

$R_\infty$ = carrying capacity of *R* in the medium

$k_{lys}$ = rate coefficient for lysing of *R* by LysisE

*Lys(t)* =concentration of LysisE at time *t*

The $k_{lys}$ value was estimated using values from McCready et al. (McCready et al., 2019).

The AI-1 induced expression of LasR and subsequently LysisE in the receiver cells was modeled based on the models developed by Sams lab (Claussen et al., 2013; Welch et al., 2020).

We made the following assumption regarding concentration of *lasR* transcript.

$$\frac{dlasR(t)}{dt} = 2\frac{dR(t)}{dt} \qquad (4)$$

Where

$lasR(t)$= concentration of *lasR* at time *t*, $lasR(0) = 2R(0)$

For the $H_2O_2$ induced expression of *lasI* and subsequently production of AI-1, we used Michaelis-Menten kinetics.

$$\frac{dH_2O_2(t)}{dt} = -\frac{k_1 H_2O_2(t) 2T(t)}{k_{mH_2O_2} + 2T(t)}$$

Where

$H_2O_2(t)$ = concentration of $H_2O_2$ at time $t$

$k_1$= rate coefficient for interaction between $H_2O_2$ and OxyR

We made the assumption that $OxyR(t) \sim 2T(t)$

$k_{mH_2O_2}$= Michaelis constant for interaction between $H_2O_2$ and OxyR

$$\frac{dAI1RBS(t)}{dt} = \frac{k_1 H_2O_2(t) 2T(t)}{k_{mH_2O_2} + 2T(t)} - k_{-AI1RBS} AI1RBS(t) \qquad (5)$$

Where

$AI1RBS(t)$ =concentration of *AI1RBS* complex at time $t$

$k_{-AI1RBS}$= *AI1RBS* degradation rate coefficient

The remaining equations are:

$$\frac{dAI1(t)}{dt} = k_{fAI1} AI1RBS(t) - 2k_{3+} LasR_2(t) AI1(t) + k_{3-} LasR_2AI1(t) + 2k_{4-} LasR_2AI1_2(t) - k_{4+} LasR_2AI1(t) AI1(t) \qquad (6)$$

Where

$AI1(t)$ =concentration of *AI*1 at time $t$

$LasR_2AI1(t)$ =concentration of *LasR* dimer with 1 *AI*1 molecule at time $t$

$LasR_2AI1_2(t)$ = concentration of *LasR* dimer with 2 *AI*1 molecules at time $t$

$k_{fAI1}$= rate coefficient for formation of *AI*1

$k_{3\pm}$ = Formation/dissociation rate coefficients for formation of $LasR_2AI1$

$k_{4\pm}$ = Formation/dissociation rate coefficients for formation of $LasR_2AI1_2$

$$\frac{dLasR(t)}{dt} = k_{fLasR} lasR(t) + 2k_{2-} LasR_2(t) - k_{-LasR} LasR(t) - 2k_{2+} LasR(t)^2 \quad (7)$$

Where

$LasR(t)$ =concentration of *LasR* monomer at time *t*

$LasR_2(t)$ = concentration of *LasR* dimer at time *t*

$k_{fLasR}$=formation rate coefficient for *LasR* monomer

$k_{2\pm}$ = Formation/dissociation rate coefficients for formation of $LasR_2$

$k_{-LasR}$ =*LasR* monomer degradation coefficient

$$\frac{dLasR_2(t)}{dt} = k_{2+}LasR(t)^2 - k_{2-}LasR_2(t) + k_{3-}LasR_2AI1(t) - 2k_{3+}LasR_2(t)AI1(t) \quad (8)$$

$$\frac{dLasR_2AI1(t)}{dt} = 2k_{3+}LasR_2(t)AI1(t) - k_{3-}LasR_2AI1(t) + 2k_{4-}LasR_2AI1_2(t) - k_{4+}LasR_2AI1(t)AI1(t) \quad (9)$$

$$\frac{dLasR_2AI1_2(t)}{dt} = k_{4+}LasR_2AI1(t)AI1(t) - 2k_{4-}LasR_2AI1_2(t) \quad (10)$$

$$\frac{dPLux(t)}{dt} = 2\frac{dR(t)}{dt} + k_{-PLux_a}PLux_a(t) - \frac{k_5PLux(t)LasR_2AI1_2(t)}{k_{mLasR_2AI1_2}+LasR_2AI1_2(t)} \quad (11)$$

Where

$PLux(t)$ = concentration of *PLux* promoter at time *t*, $PLux(0) = 2R(0)$

$PLux_a(t)$ =concentration of active *PLux* promoter at time *t*

$k_5$= rate coefficient for interaction between $LasR_2AI1_2$ and *PLux*

$k_{-PLux_a}$ =off rate coefficient for active *PLux* promoter

$k_{mLasR_2AI1_2}$= Michaelis constant for interaction between *PLux* promoter and $LasR_2AI1_2$complex

$$\frac{dPLux_a(t)}{dt} = \frac{k_5PLux(t)LasR_2AI1_2(t)}{k_{mLasR_2AI1_2(t)}+LasR_2AI1_2(t)} - k_{-PLux_a}PLux_a(t) \quad (12)$$

$$\frac{dLys(t)}{dt} = k_{fLys1}PLux_a(t) - k_{-Lys}Lys(t) \quad (13)$$

Where

$k_{fLys1}$ = rate coefficient for production of *LysisE* using *PLux* promoter

$k_{-Lys}$= degradation rate coefficient for *LysisE*

While some of the parameters (e.g., $k_{fLasR}$, $k_{-LasR}$) are available from the published models (Claussen et al., 2013; Welch et al., 2020) and references therein, for the unknowns parameters, we set initial values based on values from similar systems and intuition. We then conducted a sensitivity analysis for each unknown parameter and found which ones had the highest control on the behavior of the system. We then adjusted those values, in the order of highest sensitivity to lowest, with the aim of minimizing SSE between model predictions and experimental measurements. The most sensitive values in descending order were $k_1$, $k_{fAI1}$, and $k_{-AI1RBS}$. For the OxyR-LasI-LAA, K31-LasI-LAA, and K33-LasI-LAA constructs in Figure 2, we needed to optimize these three most sensitive values, all related to binding of $H_2O_2$, to achieve the optimum agreement between model predictions and experimental measurements. This agrees with our expectation of how introductions of these construct would change the dynamics of the system. The sets of values for the three constructs can be found in Table S2.

For electrogenetic induction of transmitter-receiver co-cultures (Figure 3), we used the set of values from $H_2O_2$ addition experiments with the OxyR-LasI-LAA construct. We then for each set of measurements varied the initial concentration of $H_2O_2$ to get optimal agreement between measurements and model predictions. The predicted "apparent $H_2O_2$" concentrations are shown in Figure 3E.

For the mutations of the S129 codon of the LasR transcriptional activator (Figure 4A), we varied the values of $k_{2+}$, $k_{3+}$, and $k_{4+}$ to see which value had the most control over the dynamics of the system while minimizing SSE between experiments and our model predictions. We found that manipulating the values that affect binding of $H_2O_2$ to the dimer of LasR (i.e., $k_{3+}$, and $k_{4+}$) provide us with the best results. The predicted values of $k_{3+}$, and $k_{4+}$ for the mutants are recorded in Table S2.

For the ECF/antiECF systems for programming a time-delay in lysis response, we first used fluorescent measurements for LasR(S129W)-sCFP(3A) (Figure S7) and LasR-J23106-ECF-sCFP(3A) (Figure S8) to calculate the value for the rate coefficient for production of LysisE from the $P_{ECF}$ promoter ($k_{fLys2}$). The following equations were then used to describe the dynamics of ECF formation and interaction with antiECF. For each Anderson promoter variant, we used a different initial antiECF value that is listed in Table S2.

$$\frac{dECF(t)}{dt} = k_{fECF}PLux_a(t) - k_6 antiECF(t)ECF(t) - k_7 ECF(t)PEcf(t) \quad (14)$$

$$\frac{dantiECF(t)}{dt} = C_n \frac{dR(t)}{dt} - k_6 antiECF(t)ECF(t) \quad (15)$$

Where

$ECF(t)$ = concentration of *ECF* at time *t*

$antiECF(t)$ = concentration of *antiECF* at time *t*

$PEcf(t)$ = concentration of *ECF* at time *t*, $PEcf(0) = 2R(0)$

$k_{fECF}$ = production rate coefficient for *ECF*

$k_6$= rate coefficient of interaction between *ECF* and *antiECF*

$k_7$= rate coefficient of interaction between *ECF* and *PEcf*

$C_n$= copy number of antiECF molecules per cell.

We used a $C_n$ value of 2 for variant J23119 (Yan & Fong, 2017) and optimized for a value of $k_6$ which we used for all the other variants as well.

$$\frac{dPEcf(t)}{dt} = 2\frac{dR(t)}{dt} - k_7 ECF(t)PEcf(t) + k_{-pEa}pEa(t) \qquad (16)$$

Where

$pEa(t)$ = concentration of active *PEcf* at time *t*

$k_{-pEa}$ = off rate coefficient for the active *PEcf*

$$\frac{dpEa(t)}{dt} = k_7 ECF(t)PEcf(t) - k_{-pEa}pEa(t) \qquad (17)$$

$$\frac{dLys(t)}{dt} = k_{fLys2}pE_a(t) - k_{-Lys}Lys(t) \qquad (18)$$

Where

$k_{fLys2}$ = rate coefficient for production of *LysisE* from the *PEcf* promoter

For all simulations, we used the measured $OD_{600}$ values to set the initial concentrations of the receiver and transmitter cells based on the ratios reported in the manuscript.

### Supplemental Tables

#### Tables S1. List of all plasmids used in this work with relevant features

| Plasmid Name | Key Features |
|---|---|
| OxyR-LasI-LAA | pBR322, AmpR; Constitutive *oxyR* expression from $P_{tet}$; $H_2O_2$ induced *lasI(LAA)* expression (encoding LasI with a LAA SsrA degradation tag) from OxyR activated $P_{oxyS}$. Strongest transmitter. |
| K31-LasI-LAA | pBR322, AmpR; Constitutive *oxyR* expression from $P_{tet}$; $H_2O_2$ induced *lasI-laa* expression from OxyR activated $P_{katG}$ combined with *rbs31*. Intermediate transmitter. |
| K33-LasI-LAA | pBR322, AmpR; Constitutive *oxyR* expression from $P_{tet}$; $H_2O_2$ induced *lasI-laa* expression from OxyR activated $P_{katG}$ combined with *rbs33*. Weakest transmitter. |
| K31-LysisE | pBR322, KanR; Constitutive *oxyR* expression from $P_{tet}$; $H_2O_2$ induced *lysisE* expression from OxyR activated $P_{katG}$ combined with *rbs31*. Monoculture model plasmid. |
| K33-LysisE | pBR322, KanR; Constitutive *oxyR* expression from $P_{tet}$; $H_2O_2$ induced *lysisE* expression from OxyR activated $P_{katG}$ combined with *rbs33*. Monoculture model plasmid. |
| LasR(WT)-LysisE | pSC101, KanR; Constitutive *lasR(WT)* expression from $P_{J23119}$; AI-1 induced *lysisE* expression from LasR(WT) activated $P_{lux}$ (responding to sub 10 nM AI-1 concentrations). Receiver plasmid. |
| LasR(S129W)-LysisE | pSC101, KanR; Constitutive *lasR(S129W)* expression from $P_{J23119}$; AI-1 induced *lysisE* expression from LasR(S129W) activated $P_{lux}$ (AI-1 $EC_{50}$ of 77 nM). Receiver plasmid. |
| LasR(S129F)-LysisE | pSC101, KanR; Constitutive *lasR(S129F)* expression from $P_{J23119}$; AI-1 induced *lysisE* expression from LasR(S129F) activated $P_{lux}$ (AI-1 $EC_{50}$ of 177 nM). Receiver plasmid. |
| LasR(S129T)-LysisE | pSC101, KanR; Constitutive *lasR(S129T)* expression from $P_{J23119}$; AI-1 induced *lysisE* expression from LasR(S129T) activated $P_{lux}$ (AI-1 $EC_{50}$ of 870 nM). Receiver plasmid. |
| LasR(S129M)-LysisE | pSC101, KanR; Constitutive *lasR(S129M)* expression from $P_{J23119}$; AI-1 induced *lysisE* expression from LasR(S129M) activated $P_{lux}$ (AI-1 $EC_{50}$ of 6600 nM). Receiver plasmid. |
| LasR-J23106-ECF-LysisE | pSC101, KanR; Constitutive *lasR(WT)* expression from $P_{J23119}$; AI-1 induced *ecf20* expression from LasR(WT) activated $P_{lux}$; *lysisE* expression from ECF20 activated $P_{ECF}$; Constitutive expression of *anti-ECF20* from $P_{J23106}$. Weakest expression of anti-ECF. Receiver plasmid. |
| LasR-J23100-ECF-LysisE | pSC101, KanR; Constitutive *lasR(WT)* expression from $P_{J23119}$; AI-1 induced *ecf20* expression from LasR(WT) activated $P_{lux}$; *lysisE* expression from ECF20 activated $P_{ECF}$; Constitutive expression of *anti-ECF20* from $P_{J23100}$. Intermediate expression of anti-ECF. Receiver plasmid. |
| LasR-J23119-ECF-LysisE | pSC101, KanR; Constitutive *lasR(WT)* expression from $P_{J23119}$; AI-1 induced *ecf20* expression from LasR(WT) activated $P_{lux}$; *lysisE* expression from ECF20 activated $P_{ECF}$; |

| | |
|---|---|
| | Constitutive expression of *anti-ECF20* from $P_{J23119}$. Strongest expression of anti-ECF. Receiver plasmid. |
| LasR(WT)-mVenus | pSC101, KanR; Constitutive *lasR(WT)* expression from $P_{J23119}$; AI-1 induced *mVenus* expression from LasR(WT) activated $P_{lux}$ (responding to sub 10 nM AI-1 concentrations). Receiver plasmid, fluorescent reporter. |
| LasR(S129W)-sCFP(3A) | pSC101, KanR; Constitutive *lasR(S129W)* expression from $P_{J23119}$; AI-1 induced *sCFP(3A)* expression from LasR(S129W) activated $P_{lux}$ (AI-1 $EC_{50}$ of 77 nM). Receiver plasmid, fluorescent reporter. |
| LasR(S129F)-sCFP(3A) | pSC101, KanR; Constitutive *lasR(S129F)* expression from $P_{J23119}$; AI-1 induced *sCFP(3A)* expression from LasR(S129F) activated $P_{lux}$ (AI-1 $EC_{50}$ of 177 nM). Receiver plasmid, fluorescent reporter. |
| LasR(S129T)-sCFP(3A) | pSC101, KanR; Constitutive *lasR(S129T)* expression from $P_{J23119}$; AI-1 induced *sCFP(3A)* expression from LasR(S129T) activated $P_{lux}$ (AI-1 $EC_{50}$ of 870 nM). Receiver plasmid, fluorescent reporter. |
| LasR(S129M)-sCFP(3A) | pSC101, KanR; Constitutive *lasR(S129M)* expression from $P_{J23119}$; AI-1 induced *sCFP(3A)* expression from LasR(S129M) activated $P_{lux}$ (AI-1 $EC_{50}$ of 6600 nM). Receiver plasmid, fluorescent reporter. |
| LasR-J23106-ECF-sCFP(3A) | pSC101, KanR; Constitutive *lasR(WT)* expression from $P_{J23119}$; AI-1 induced *ecf20* expression from LasR(WT) activated $P_{lux}$; *sCFP(3A)* expression from ECF20 activated $P_{ECF}$; Constitutive expression of *anti-ECF20* from $P_{J23106}$. Weakest expression of anti-ECF. Receiver plasmid, fluorescent reporter. |
| LasR-J23100-ECF-sCFP(3A) | pSC101, KanR; Constitutive *lasR(WT)* expression from $P_{J23119}$; AI-1 induced *ecf20* expression from LasR(WT) activated $P_{lux}$; *sCFP(3A)* expression from ECF20 activated $P_{ECF}$; Constitutive expression of *anti-ECF20* from $P_{J23100}$. Intermediate expression of anti-ECF. Receiver plasmid, fluorescent reporter. |
| LasR-J23119-ECF-sCFP(3A) | pSC101, KanR; Constitutive *lasR(WT)* expression from $P_{J23119}$; AI-1 induced *ecf20* expression from LasR(WT) activated $P_{lux}$; *sCFP(3A)* expression from ECF20 activated $P_{ECF}$; Constitutive expression of *anti-ECF20* from $P_{J23119}$. Strongest expression of anti-ECF. Receiver plasmid, fluorescent reporter. |

**Table S2. Table of model parameters**

| Parameter | Value |
|---|---|
| for K31-LysisE | |
| $K_{i,H2O2}$<br>$T_\infty$<br>$k_T$<br>$k_{dtT}$<br>$k_{dtH_2O_2}$ | 30 μM<br>0.00182 nM<br>0.5 $hr^{-1}$<br>0.001 $hr^{-1}$<br>0.0005 $\mu M^{-1}$ $hr^{-1}$ |
| LasR(WT)-LysisE | |
| $k_R$<br>$R_\infty$<br>$k_{dtR}$<br>$k_{lys}$<br>$k_{fLasR}$<br>$k_1$<br>$k_{2-}$<br>$k_{2+}$<br>$k_{-LasR}$<br>$k_{3-}$<br>$k_{3+}$<br>$k_{4-}$<br>$k_{4+}$<br>$k_5$<br>$k_{fLys1}$<br>$k_{-Lys}$<br>$k_{fAI1}$<br>$k_{-AI1RBS}$<br>$k_{-PLuxa}$<br>$k_{mLasR2AI1}$<br>$k_{mH2O2}$ | 0.75 $hr^{-1}$<br>0.00143 nM<br>0.002 $hr^{-1}$<br>7700 $nM^{-1}$ $hr^{-1}$<br>1000 $hr^{-1}$<br>250 $nM^{-1}$ $hr^{-1}$<br>1 $hr^{-1}$<br>1390 $hr^{-1}$<br>20 $hr^{-1}$<br>72 $hr^{-1}$<br>8 $nM^{-1}$ $hr^{-1}$<br>99 $hr^{-1}$<br>11 $nM^{-1}$ $hr^{-1}$<br>100 $nM^{-1}$ $hr^{-1}$<br>1 $hr^{-1}$<br>1.2 $hr^{-1}$<br>3.5 $hr^{-1}$<br>1.5 $hr^{-1}$<br>1 $hr^{-1}$<br>2.2 nM<br>1 nM |
| K31-LasI-LAA, and K33-LasI-LAA constructs | |
| $k_1$ (RBS31)<br>$k_1$ (RBS33)<br>$k_{fAI1}$ (RBS31)<br>$k_{fAI1}$ (RBS33)<br>$k_{-AI1RBS}$ (RBS31)<br>$k_{-AI1RBS}$ (RBS33) | 187.5 $nM^{-1}$ $hr^{-1}$<br>3 $nM^{-1}$ $hr^{-1}$<br>2.6 $hr^{-1}$<br>2.4 $hr^{-1}$<br>4 $hr^{-1}$<br>0.55 $hr^{-1}$ |
| LasR S129 codon mutations | |
| $k_{3+}$ (LasR S129W)<br>$k_{3+}$ (LasR S129F)<br>$k_{3+}$ (LasR S129T)<br>$k_{4+}$ (LasR S129W)<br>$k_{4+}$ (LasR S129F)<br>$k_{4+}$ (LasR S129T) | 0.8 $nM^{-1}$ $hr^{-1}$<br>0.16 $nM^{-1}$ $hr^{-1}$<br>0.012 $nM^{-1}$ $hr^{-1}$<br>1.1 $nM^{-1}$ $hr^{-1}$<br>0.12 $nM^{-1}$ $hr^{-1}$<br>0.016 $nM^{-1}$ $hr^{-1}$ |
| ECF-antiECF system | |
| $k_6$<br>$C_n$ (J23106)<br>$C_n$ (J23100)<br>$C_n$ (J23119)<br>$k_7$<br>$k_{-pEa}$<br>$k_{fLys2}$ | 10800 $nM^{-1}$ $hr^{-1}$<br>0.003<br>1.75<br>2<br>100 $nM^{-1}$ $hr^{-1}$<br>1 $hr^{-1}$<br>24 $hr^{-1}$ |

**Supplementary Figures**

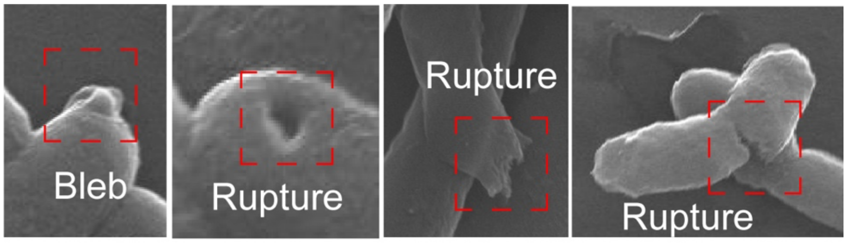


**Figure S1. Scanning Electron Microscopy (SEM) of *E. coli* cells after LysisE induction.** LysisE induced membrane blebs (left-most SEM image) and bursts (right three SEM images) shortly after expression reaches a critical threshold.

A.)

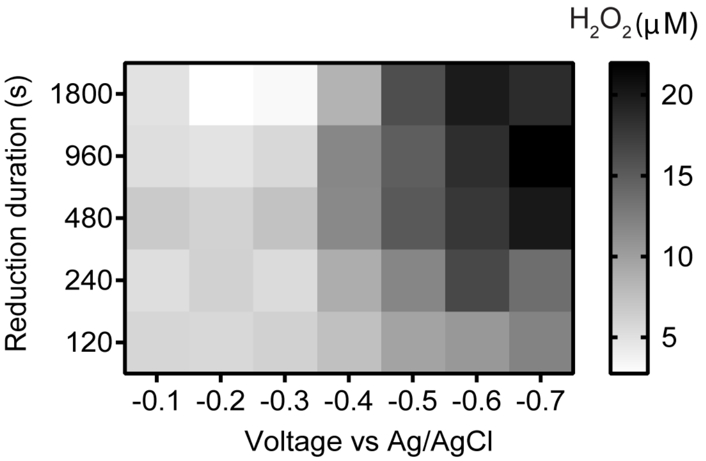


B.)

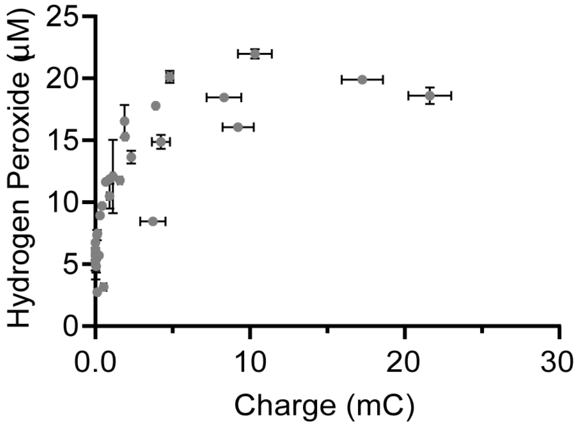


**Figure S2. Electrogenetic production of hydrogen peroxide in a 3-electrode chamber without cells.** Hydrogen peroxide concentrations (measured via colorimetric assays) mapped to program inputs (**A**) and total charge (**B**). All experiments were performed in triplicates.

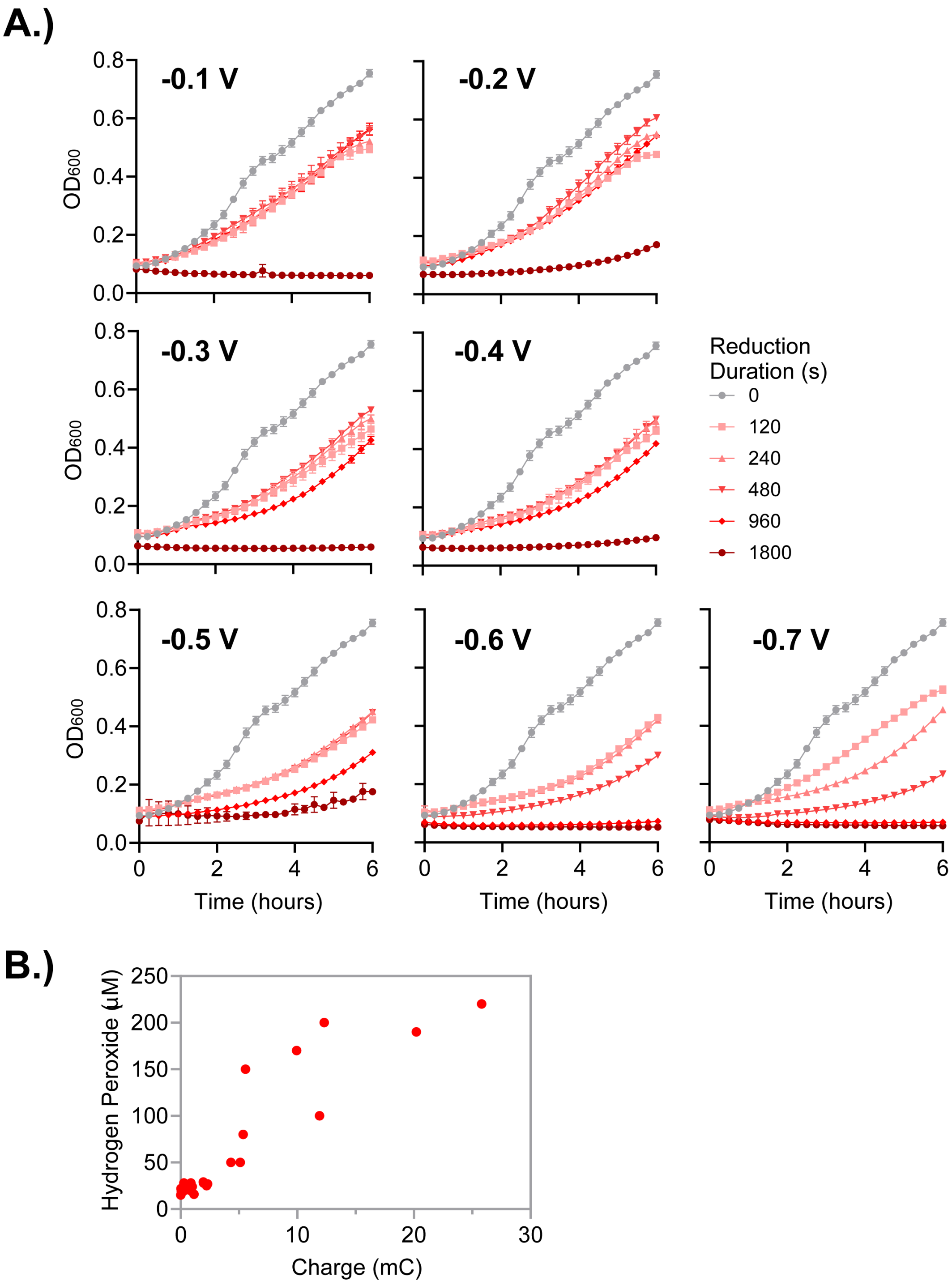


**Figure S3. Electrogenetic lysis of the K31-LysisE construct. A.)** Monocultures were induced with the indicated time and voltage at 0.1 $OD_{600}$ and were monitored for 6 hours. Growth curves post induction are shown. **B.)** Apparent initial $H_2O_2$ values from model predictions versus charge. In general, locally experienced hydrogen peroxide was significantly greater than the amount actually produced. All experiments were performed in triplicates.

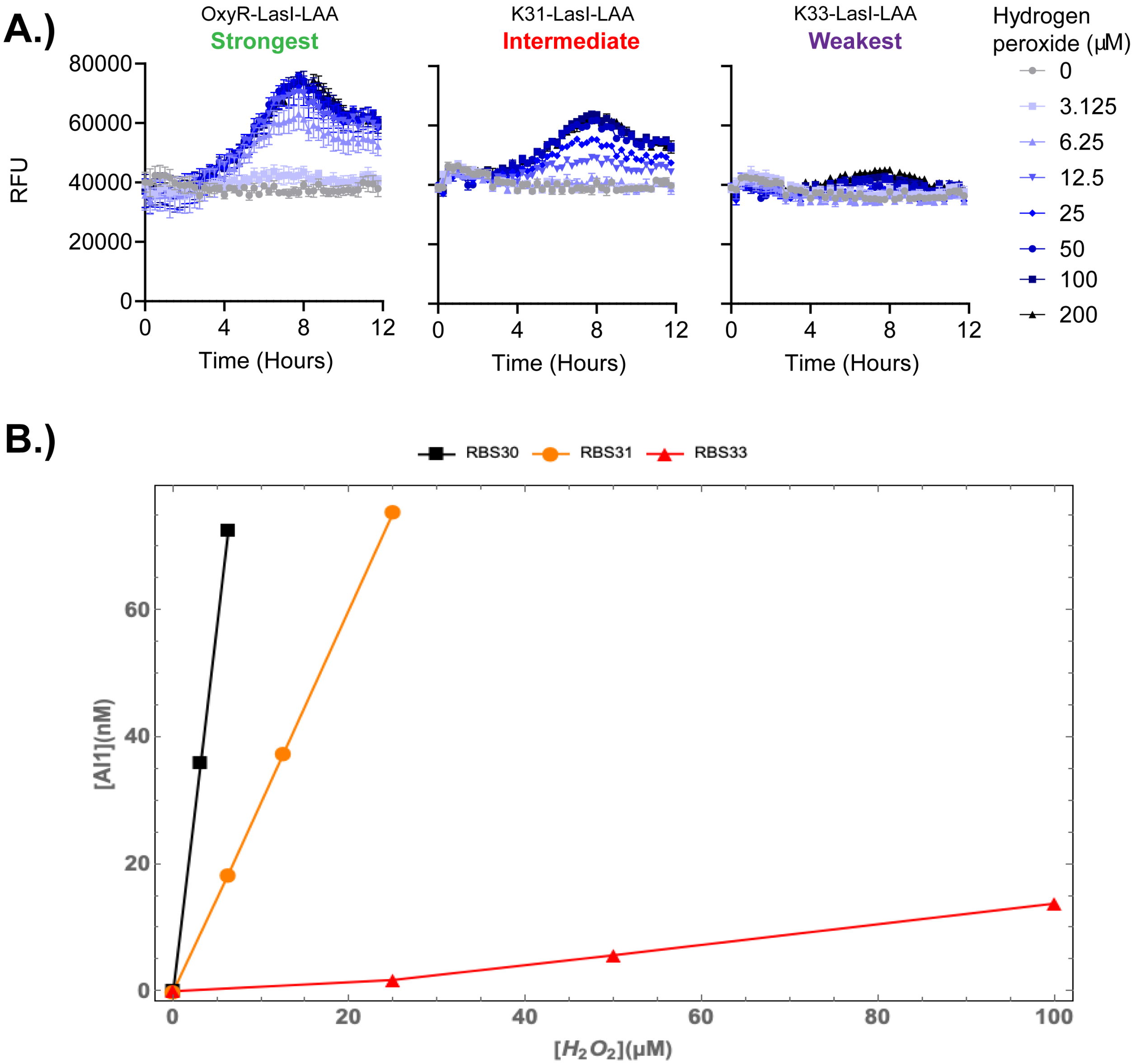


**Figure S4. Altering the lysis response through transmitter design.** A.) Fluorescent response of transmitter-receiver cocultures initially containing LasR(WT)-mVenus receiver cells and 1% transmitter cells, either OxyR-LasI-LAA (green, left), K31-LasI-LAA (red, middle), or K33-LasI-LAA (purple, right) that are responsive to different $H_2O_2$ levels. Cells were induced (at time 0) with the indicated $H_2O_2$ concentration (between 0-200 μM). All experiments were performed in triplicates. B.) Model prediction of AI-1 level after 4 h. RBS30 is OxyS-LasI-LAA, RBS31 is K31-LasI-LAA, and RBS33 is K33-LasI-LAA.

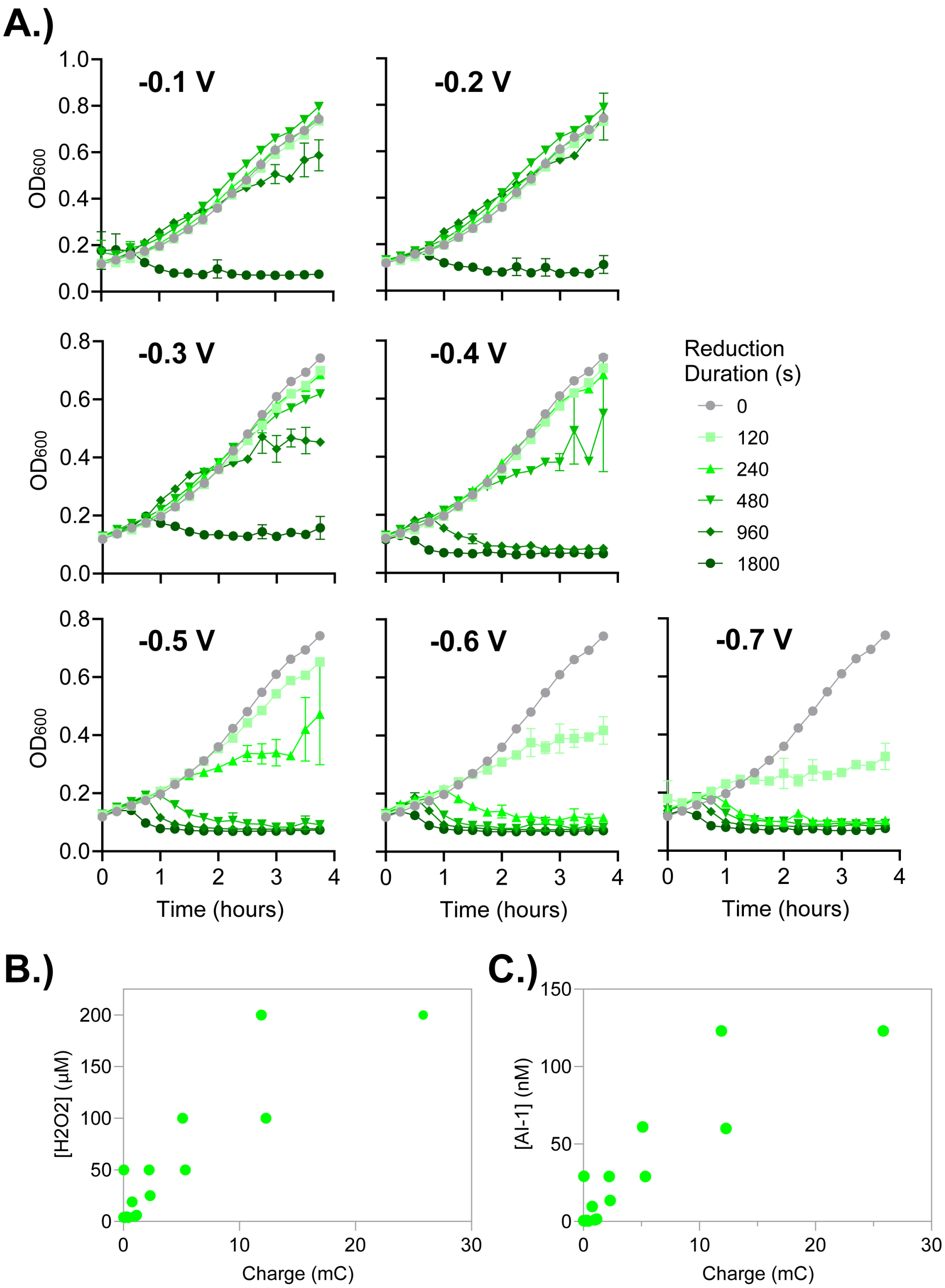


**Figure S5. Electrogenetic induction of the transmitter-receiver lysis system. A.)** Cocultures of OxyR-LasI-LAA transmitters and LasR(WT) receivers were induced with the indicated time and voltage at a total $OD_{600}$ of 0.1 and a transmitter density of 1% and were monitored for 4 hours. Growth curves post induction are shown. **B-C.)** Model fitted predictions of apparent initial hydrogen peroxide (B) and AI-1 (C) versus the input charge. All experiments were performed in triplicates.

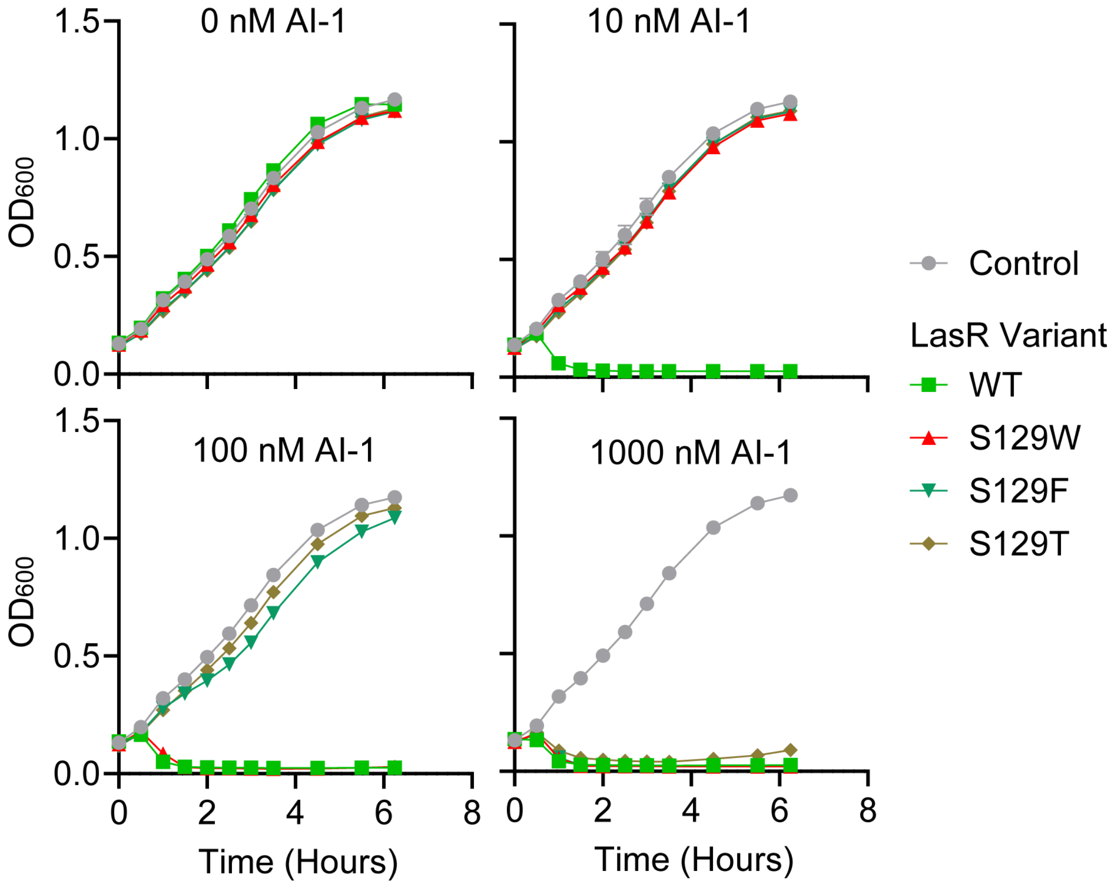


**Figure S6. AI-1 induction of LysisE receiver cells containing LasR S129 variants.** Monocultures of cells at 0.1 $OD_{600}$ were induced with 3-oxo-C12-HSL (i.e. AI-1) at the indicated concentrations. Growth post-induction was monitored for 6 h by $OD_{600}$.

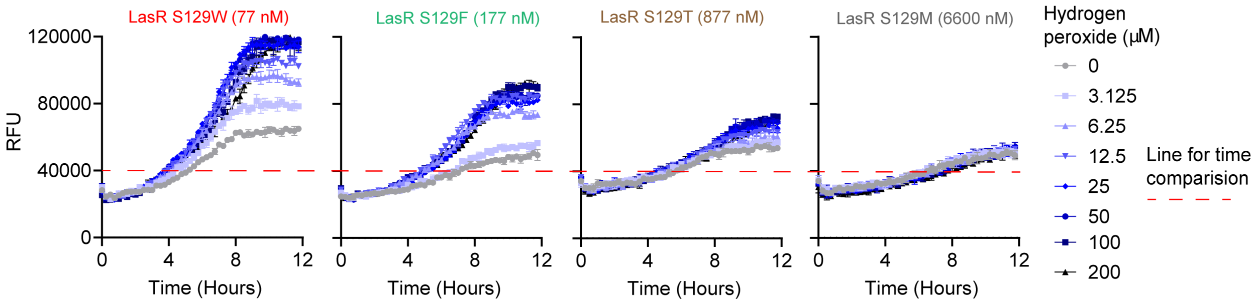


**Figure S7. Fluorescence response from the LasR variant transmitter-receiver systems using fluorescent reporters.** Cocultures initially containing 1% OxyR-LasI-LAA transmitters and indicated LasR variant receiver constructs controlling expression of *sCFP(3A)* (replacing *lysisE*). Cells were induced (at time 0) with the indicated $H_2O_2$ concentration (between 0-200 μM) and fluorescence was monitored for 12 h post-induction. The defining circuit feature is labeled above each plot. A red-dashed line is drawn to show the relative delay of each system to reach a set threshold. All experiments were performed in triplicates.

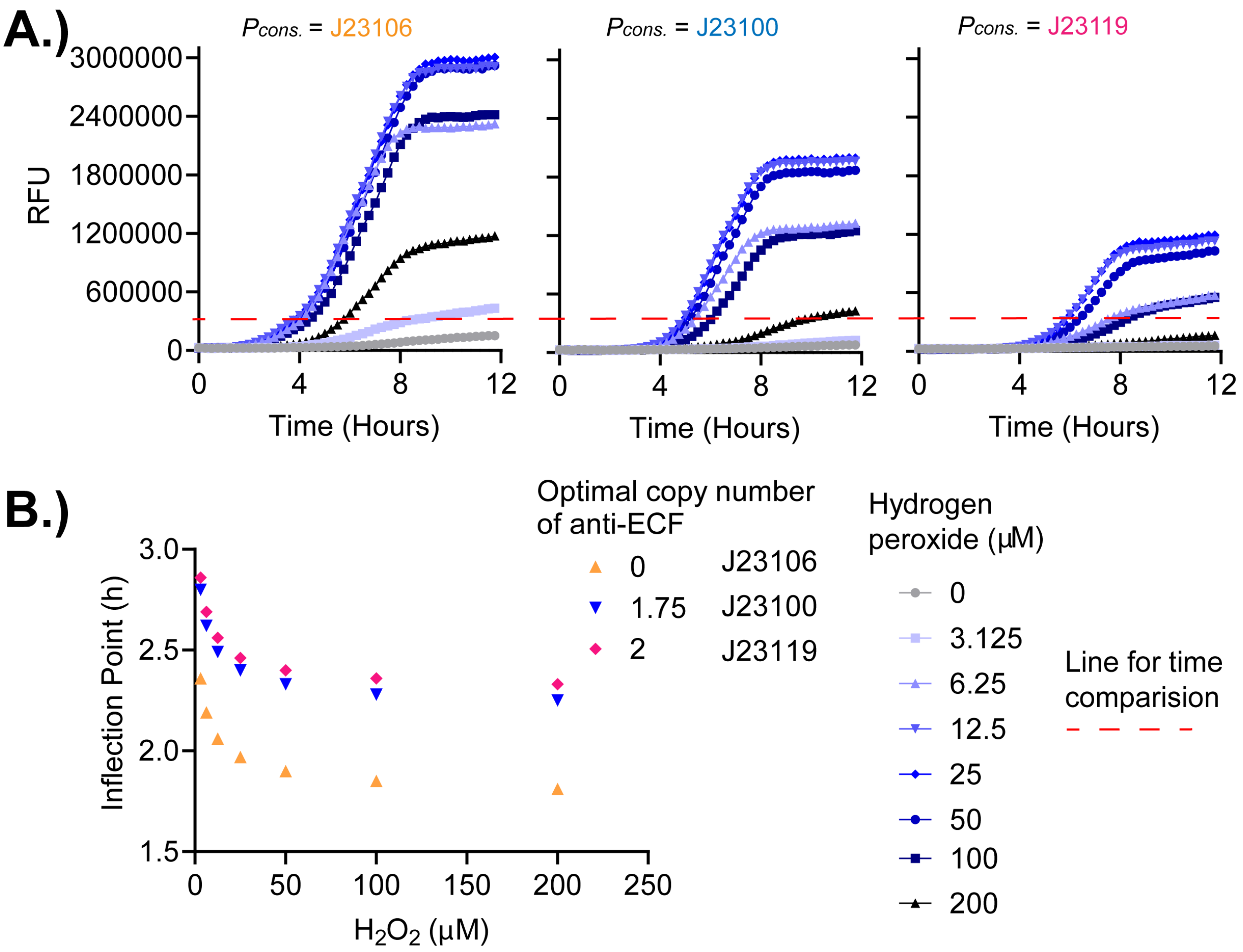


**Figure S8. Fluorescence response from the ECF/anti-ECF transmitter-receiver systems using fluorescent reporters. A.)** Cocultures initially containing 1% OxyR-LasI-LAA transmitters and indicated ECF/anti-ECF receiver constructs controlling expression of *sCFP(3A)* (replacing *lysisE*). Cells were induced (at time 0) with the indicated $H_2O_2$ concentration (between 0-200 µM) and fluorescence was monitored for 12 h post-induction. The defining circuit feature is labeled above each plot. A red-dashed line is drawn to show the relative delay of each system to reach a set threshold. **B.)** The model predicted inflection points of each system as a function of hydrogen peroxide. The optimal copy number for matching the lysis data in Figure 5 and fluorescent data in Figure S8 is shown for each variant (J23106 = 0; J23100 = 1.75, J23119 = 2).

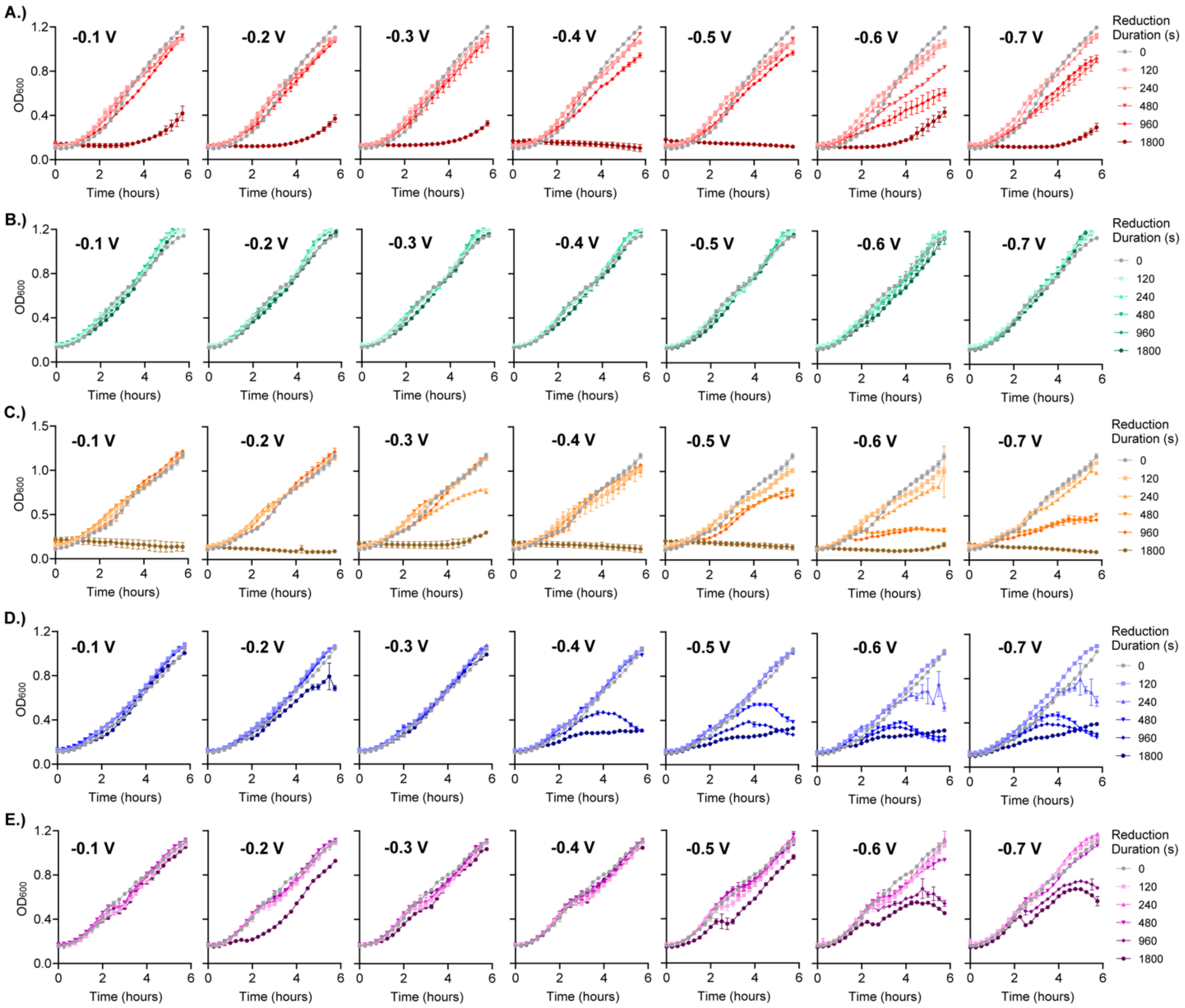


**Figure S9. Electrogenetic induction of time-delay systems with transmitter-receiver cocultures.** Cocultures were induced with the indicated time and voltage at a total $OD_{600}$ of 0.1 and a OxyR-LasI-LAA transmitter density of 1% and were monitored for 6 hours. Growth curves post induction are shown. The receiver variants are as follows: **A.)** LasR(S129W), **B.)** LasR(S129F), **C.)** J23106, **D.)** J23100, **E.)** J23119. All experiments were performed in triplicates.

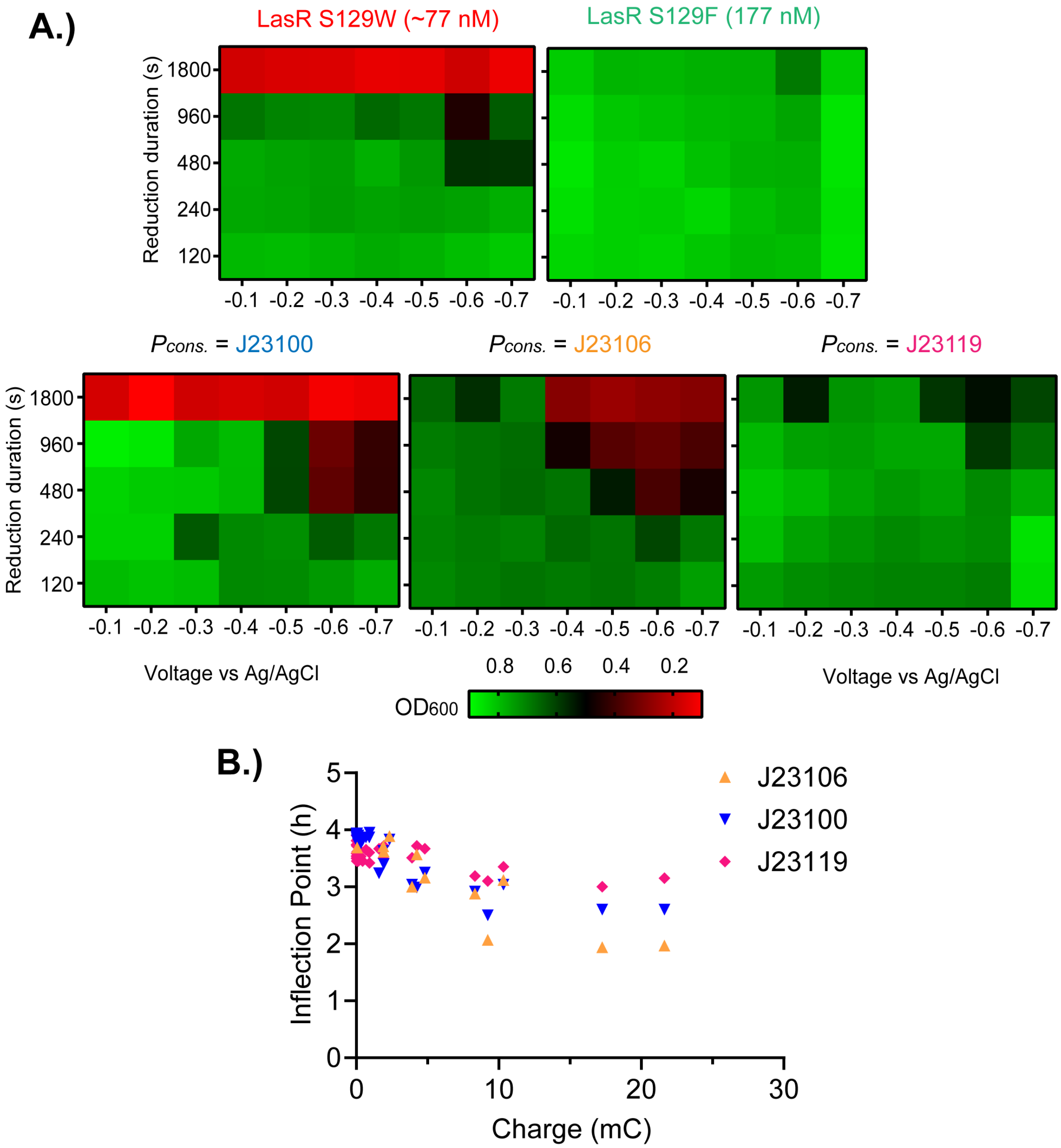


**Figure S10. Analyses of electrogenetic lysis of LasR variant and anti-ECF/ECF variant transmitter-receiver systems. A.)** Heatmap shows the final $OD_{600}$ after 4 h of growth in Figure S9 as functions of voltage inputs (horizontal-axis) and reduction durations (vertical-axis). Energy inputs increase towards the upper-right corner of the heatmap. **B.)** The model predicted inflection points versus charge. Copy numbers effectively capture observed trends.
